# Statistical knockoffs improve biomarker discovery from transcriptomic data

**DOI:** 10.1101/2025.07.04.663147

**Authors:** Julie Cartier, Johanna Lagoas, Youmna Ayadi, Adeline Fermanian, Chloé-Agathe Azencott, Florian Massip

**Affiliations:** Centre for Computational Biology, MinesParis, PSL University, 60 bd Saint-Michel, 75272, Paris, France; Institut Curie, PSL University, 11 rue Pierre et Marie Curie, 75005, Paris, France; U1331, INSERM, 11 rue Pierre et Marie Curie, 75005, Paris, France; LOPF, LOPF Califrais’Machine Learning Lab, Paris, France

**Keywords:** Knockoffs, Variable selection, Transcriptomic

## Abstract

Advances in sequencing technologies have enabled the generation of large amounts of data, offering new possibilities to identify relationships between biological units (*e.g*. genes) and phenotypic traits (*e.g*. disease outcomes). Yet, identifying these associations using variable selection methods remains challenging due to the high dimension (*p* ≫ *n*) and the correlation structure of the data. To address these challenges, we study the applicability of the knockoff (KO) procedure. Introduced by Barber and Candès in 2015, the KO variable selection procedure has shown promising results on real biological data, such as Genome Wide Association Studies. This method seeks to identify the truly important predictors by overcoming the correlation structure between variables while controlling the false discovery rate. Here, we study the applicability of the KO procedure on transcriptomic data in a classification setting. We conduct an extensive simulation study using real transcriptomic data to evaluate the performance of the KO framework in the context of high-dimensional classification. We find that the KO framework outperforms widely used variable selection models, and that using KO aggregation to mitigate the effect of KO stochasticity improves stability while maintaining the same power. Finally applied to three real transcriptomic datasets, the KO framework made very few discoveries, highlighting its conservative nature and suggesting that other methods may substantially overestimate the number of relevant features.

## 1 Introduction

Thanks to the progress of sequencing technologies, it is now possible to collect and access sequencing and genomic information for large cohorts of patients, paving the way for the use of these technologies for many biotechnological and medical applications. In particular, patient stratification and disease grading based on the results of a sequencing experiment is seen as a promising avenue for several diseases, and in cancer in particular [1, 2]. But despite a lot of research on the topic, discovering biomarkers using transcriptomic data remains challenging. In particular, it has often been noted that transcriptomic signatures for a disease are not always reliable [3], difficult to reproduce, and that the genes selected as biomarkers are usually highly unstable [4, 5].

Several reasons make biomarker discovery particularly difficult in this context. First, despite the increase in cohort sizes, the number of explanatory variables remains much larger than the number of patients, which is well known to be a difficult statistical setup (often called “high dimension” or “large *p* small *n*”) [6, 7]. An additional difficulty arises due to the high level of correlation between the different gene’s expression. Indeed, genes that interact to take part to common biological functions are often co-regulated by similar mechanisms, inducing correlation between genes’ expressions. As a result, it becomes difficult to differentiate genes that are truly associated to the outcome of interest from those that appear associated to the outcome solely because they are correlated to a causal gene.

Recently, a novel statistical method named Knockoff (KO) [8] has been proposed to tackle the problem of variable selection for correlated features in high dimension. KO have been applied to a range of biological questions, in particular in Genome-Wide Association Studies [9, 10, 11], where it was demonstrated that KO allowed to control the number of false discoveries. However, up to now, only few studies have suggested to apply KO to transcriptomic data [12, 13, 14] and to the best of our knowledge, applying KO to transcriptomic data, in high dimension, for classification, remains unexplored. Hence, whether KO can improve variable selection for transcriptomic data remains to be studied. In addition, because KO is a flexible method, identifying the optimal way to apply it to real data is complex, which can prevent its use in practice. For instance, many methods have been developed to generate KO matrices whose power might depend on the specificities of users’ data.

Moreover, one particular difficulty with biomarker discovery is that one often obtains unstable results, *i.e*. features selected by the model tend to change a lot with small perturbations of the input data, making it especially hard to transfer results from one cohort of patients to the next. While using KO has been shown to improve false discovery control, the impact of this statistical method on the stability of the selected features is rarely studied. Indeed, as KO generation is non-deterministic, results may vary between several runs of the same method. One way to alleviate this issue is to run the KO framework multiple times and aggregate the results [15, 16, 17, 14].

The goal of this paper is to study whether KO improves variable selection when applied to transcriptomic data for classification tasks. First, we perform a systematic comparison of the performance of different KO generation methods and statistics. We then conduct simulations to compare the KO framework with baselines, namely Lasso regression and Wilcoxon rank-sum tests. We also include KO aggregation in this experiment to assess its performance. Next, we evaluate the stability of the KO framework for variable selection. We focus on different selection scenarios and demonstrate that feature correlation reduces the power of the method. Finally, we apply our resulting KO method on three real-world datasets: two include lung cancer patients and one includes breast cancer patients.

### Overview of the KO framework

In this paper, KO refers to the versatile Model-X Knockoffs [18] which we briefly present here. A more detailed summary of the KO procedure can be found in the appendix (see Supp. Mat. A).

Let **y** ∈ {0,1}^*n*^, **y** = (*y*_1_, … *y*_*n*_)^*T*^ be the random variables representing the outcomes and *X =* (*X*_.,1_, …, *X*_.,*p*_) be the matrix of features (here, genes’ expressions) such that for any *j* ∈ {1, … *p*}, *X*_.,*j*_ = (*X*_1,*j*_, …, *X*_*n*,*j*_)^*T*^ ∈ ℝ^*n*^. In the following *X*_.,*j*_ will be noted *X*_*j*_. Let ℋ_0_ denote the set of indices of null features such that for all *j* ∈ ℋ_0_, **y** ⫫ *X*_*j*_ |*X*_−*j*_ where *X*_−*j*_ is the matrix *X* without the *j*^th^ column. The goal of variable selection is to identify the set of non-null features given the observations *X* and **y**, *i.e*., the set 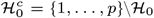. The KO framework does so while ensuring that the false discovery rate (FDR) is controlled. Therefore, running the KO framework results in the selection of a set of variables 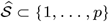 such that:

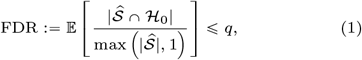

where *q* is the target FDR level.

The KO framework relies on the construction of false copies of every feature in the data matrix *X*, which will be used as control variables. These copies are gathered to form the KO matrix 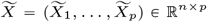. To guarantee FDR control (that is, Eq. (1)), the KO matrix needs to fulfill two conditions, namely that the distribution of the fake variables is as similar as possible to the distribution of the real features (“swap property”), and that the KO matrix 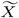 is independent of the outcomes **y** conditionally on the real feature matrix *X* (“conditional independence property”).

A statistic *W*_*j*_ is then constructed for every feature *j* based on the 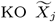 and the true feature *X*_*j*_. A high value of *W*_*j*_ must provide evidence that feature *j* is non-null. Candès et al. [18] have demonstrated that, under some assumptions, the set of features 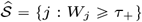, with

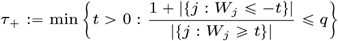

allows to control the FDR.

Additionally to control the FDR, a good variable selection method must ensure good power, that is, must be able to retrieve the non-null features. Mathematically, the power is defined by

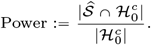

Unlike FDR control, the KO methodology does not guarantee good power by construction and power analysis is theoretically very challenging [19]. Intuitively, to have good power, 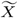 should be as different from the original matrix *X* as possible (the extreme choice 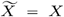 being a valid choice in terms of FDR control but having a very bad power).

In summary, the task is to build KO matrices and statistics that satisfy the swap and conditional independence properties in order to control the number of false discoveries while achieving the best possible power. Various methods have been proposed to address this problem. Here, we present a few methods that can be used in the context of high-dimensional data and classification.

### Generation of KO features

The most common methods assume that the feature matrix *X* follows a Gaussian distribution. Under this assumption, a parametrized family of distributions of 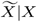 can be computed, from which a valid KO for the original features can be sampled [18]. Several families of approaches have been proposed under this assumption.

In the family of Mean Absolute Correlation (MAC) Gaussian methods, the KO features are computed to minimize the correlation between a feature *X*_*j*_ and its knockoff 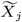 [18]. Among the different MAC approaches, the semidefinite program (SDP) is expected to be the most powerful as *p* becomes large. The MAC approaches suffer from a tendency to identify some KO of non-null original features as important features in the model instead of the non-null features themselves [20]. To address this issue, a new family of Gaussian KO generation methods called Minimum Reconstructability (MRC) has been developed. One typical approach is to maximize the conditional variance 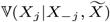, for any *j* ∈ {1, …, *p*}. This approach is known as the minimum variance-based reconstructability (MVR) method. Additionnally, conditional independent (CI) KO [19, 21] satisfies conditional independence between a feature *X*_*j*_ and its knockoff 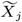, given the other covariates *X*_−*j*_.

Finally, the “Sequential Conditional Independant Pairs” (SCIP) algorithms [18] allow for the construction of valid KO with no further assumption on the distribution of the features *X*. In practice, a linear empirical version of this algorithm is often used, called the Linear Sequential Conditional Independant Pairs algorithm (LSCIP) [14, 22].

We stress that this list of methods for computing valid KO is not exhaustive. We have focused on methods particularly suitable for transcriptomic-like data. For instance, methods using deep learning models have been proposed [23, 24], but will not be considered here since our data sets have a small sample size.

### KO test statistics

Most KO statistics are built as differences between variable importance metrics. More precisely, they are obtained by applying a machine learning model to 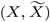 with target **y**, choosing a variable importance metric associated to the model, and computing the difference between the importance measure assigned to a feature and that assigned to its KO.

For linear settings, an intuitive option is to use the absolute values of the coefficients of a penalized logistic regression (LASSO Coefficient Difference (LCD) or Elastic-Net penalized logistic regression Coefficient Difference (EN-CD)) [8, 18] as the variable importance. Likewise, variable importance in nonlinear models can be used. These include differences in Gini impurity in random forests [20], risk reduction in boosting (RRB) in boosted trees [12], and several other boosted tree-based statistics [13], as well as approaches based on deep learning [25].

Finally, some statistics proposed in the literature cannot be directly expressed as a variable difference. One example is the masked likelihood ratio (MLR) [26], which is based on the estimation of log-likelihood ratios. This method accounts for the high correlation between a feature and its KO by lowering the absolute value of their statistic when the correlation is large. Another example is the Lasso Signed Max (LSM) [8].

### KO aggregation

The stochastic nature of KO generation methods means that applying the same KO construction method twice to the same data matrix will generate different KO matrices. This results in different subsets of selected variables at each iteration. KO aggregation addresses this issue by applying the KO procedure multiple times and “aggregating knowledge” from these runs to select variables. Several aggregation methods have been proposed, the most recent of which is the Knockoff-*π* (KOPI) [14] framework.

## Materials and Methods

To evaluate the performance of the KO framework on highdimensional transcriptomic data, we use simulated outcomes computed from real transcriptomic data. This allows us to keep the correlation structure of biological data (and its specificities such as non-Gaussianity) while knowing the true variables to recover.

### Real data sets

The CRUKPAP data [27] is composed of healthy volunteers and patients with benign lung condition (*n* = 160) and subjects diagnosed with lung cancer (*n* = 199). For all 369 samples, we have transcriptomic data from nasal epithelium samples (RNA-sequencing, 18, 072 genes) from which 749 genes have been identified as involved in patients’ response to smoke. In the following, we keep only those 749 risk genes to form the data matrix (*X*_CRUKPAP_)

The AEGIS (Airway Epithelial Gene Expression in the Diagnosis of Lung Cancer) data [28] comes from a clinical trial conducted in the USA, Canada and Europe. This data set is composed of 505 clinical patients including 308 patients diagnosed with lung cancer for whom we have gene expression levels (microarrays) of nasal epithelium (16, 127 genes). We use the set of 535 genes that were found differentially expressed in a subset of the entire cohort. For the experiments with real outcomes, we use cancer status as the outcome for both the CRUKPAP and AEGIS data sets.

Finally, the breast cancer (BC) cohort is composed of gene expression data (RNA sequencing, 18, 802 genes) from tumors of patients in the SCAN-B project (Sweden data canceronome analysis network breast initiative) [29]. These data were used to predict biomarkers in breast cancer, identifying 869 genes with non-zero weights in at least one of five biomarker classifiers. We focus on the estrogen receptor (ER), progesterone receptor (PgR) and human epidermal growth factor receptor 2 (HER2) status. Thus, our data matrix includes expression levels of these 869 genes for 2,827 samples with no missing data concerning the ER, PgR, or HER2 status. ER and PgR status described protein content, with, respectively 2, 600 and 2, 458 positives, while HER2 status reflects gene amplification, with 338 amplified cases.

We use the same pre-processing steps (quality check, batch effect correction, normalization) for these three transcriptomic data sets as in previous studies [28, 29, 27]. In all three studies mentioned above, a gene pre-filtering step based on prior knowledge or gene expression differential analysis was conducted before applying variable selection techniques, in order to reduce the dimensionality (and thus the difficulty) of the problem. Unless otherwise specified, we applied our analyses to pre-filtered data in order to compare the KO framework to the original studies.

### Outcome simulation

Unless otherwise specified, the features are standardized to have zero-mean and unit-variance, that is, for *j* ∈ {1, …, *p*}, we apply the operation 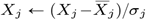, where 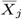 is the empirical mean of feature *j* and *σ*_*j*_ its standard deviation.

We consider two simulation settings. For each setting, we denote by *k* the number of non-zero features, that is, 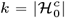. and set *k* = 10 in all experiments. We construct a vector ***β*** ∈ ℝ^*p*^ by randomly sampling *k*^′^ indices from {1, …, *d*} to which we assign values drawn uniformly from {−1, 1}, and set all remaining components of ***β*** to zero. We then multiply ***β*** with a magnitude parameter *a*.

In the linear simulation setting, we set *k* = *k*^′^ and *d* = *p*. For each *i* ∈ {1, …, *n*}, we generate the *i*^th^ component of the outcome vector **y** ∈ {0, 1}^*n*^ according to the following generalized linear model:

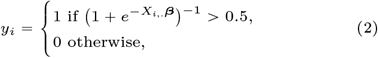

where *X*_*i*,._ is the *i*^th^ row of *X*. We take *a* = 10, calibrated such that the two classes are well separated for all data sets (see Supp. Mat. B.1: Fig. S1).

In the interaction simulation setting, we assume that *p* and *k* are even and define a new 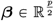 that has 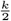 non zero components — that is 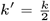 and 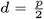. Then, the *i*^th^ component of the outcome vector **y** is defined as follows:

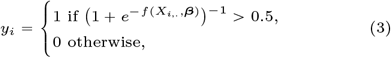

where 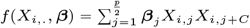 where *β*_*j*_ is the *j* cordinate of ***β*** and C is chosen in {−*j* + 1, …, −1, 1, …, *p* − *j*} such that no feature appears twice. This allows to model pairwise/quadratic interactions between variables. Note that obtaining well-separated classes in the interaction simulation setting is more difficult than in the linear setting. We set *a* to 100 and select outcome vectors that yields classes of similar proportions by trial and error (see Supp. Mat. B.1: Fig. S2).

### Implementation details

We generate KO matrices and statistics using either the Python package *knockpy* [20] (v1.3.0) or the R package *knockoff* [18] (v0.3.6) except for the LSCIP method and the EN-CD statistic, for which we use the R package *glmnet* [30] (v4.1-7). The parameter *λ*_min_ refers to the penalization obtained by minimizing the mean cross-validated error which is by default computed with the deviance (difference of log-likelihoods) error over a 10-fold cross-validation. The parameter *λ*_oracle_ is chosen *a posteriori* and corresponds to the penalization term that maximizes the ratio between true discoveries and false discoveries in a grid of values ranging from 0 to 1 in increments of 0.01. Variable selection with KOPI is performed using the code provided by [14]. The Wilcoxon rank-sum test is applied on already preprocessed data as in [31] with the R *stats* package (v.4.2.0). We correct for multiple hypotheses testing with the Benjamini-Hochberg (BH) [32] procedure. Stability selection with LASSO-penalized logistic regression is implemented with the R package *stabs* (v0.6-4). Two sampling schemes were considered: the original method of Meinhausen and Bühlmann [33] (MB) and the complementary pairs approach proposed by Shah and Samworth [34] (SS). Details are given in supplementary materials (see Supp. Mat. B.2).

### Experimental setups

We denote by *M* the number of outcome vectors simulated for each experiment. The target FDR level (*q*) sequence, used by KO-based methods, varies from 0 to 1 in increments of 0.01. For all experiments, unless otherwise specified, we use *X*_CRUKPAP_ ∈ ℝ^369×749^ as data matrix to compute **y**.

#### KO generation methods and statistics benchmark

*M* is set to 100 in the linear simulation setting and to 10 in the interaction simulation setting. From one simulation to the next, only the identity of the genes that are selected to compute **y** changes. We compare methods and statistics that seem particularly appropriate for transcriptomic-like data. We include in the comparison the SDP method as the gold standard for KO generation methods, the MVR method, as representative of MRC methods, CI-based KO construction, and the LSCIP algorithm (see Supp. Mat. Algo. 1). We use the LCD statistic for the KO generation methods benchmark since it is the gold standard statistic in the linear simulation setting.

Concerning statistics, we benchmark the LCD, since it outperforms other LASSO-based statistics (see Supp. Mat. C.1.1: Fig. S4). We also consider for linear statistics: the Elasticnet coefficient difference (EN-CD), and the MLR statistic which has shown better performance than the LCD statistic in low-dimensional linear logistic regression [26]. For tree-based methods, the results (see Supp. Fig. S5) show that none of the statistics are particularly effective, but RRB and Gini variable importance seem to be slightly more powerful. However, as RRB is extremely time-consuming to compute, we chose to use Gini-based variable importance. Given the poor performance of deep learning based statistics (see Supp. Fig. S6), we did not include them in the comparison.

#### Comparison between KO, LASSO, Wilcoxon rank-sum test, KOPI and Stability Selection

For *M* = 100 outcome vectors, we run the KO as well as the KOPI procedures with LSCIP and LCD methods. We compare it with the Wilcoxon rank-sum test combined with the BH procedure for target FDR levels in {10^−10^, 5 × 10^−10^, 10^−8^, 5 × 10^−8^, …, 0.01, 0.05, 0.5, 1}—this is the equivalent to *q* in the KO framework. We also add in the comparison a LASSO-penalized logistic regression with *λ* ranging from 0 to 0.4 in steps of 0.004, *λ*_min_ and *λ*_oracle_, and stability selection with the MB and SS sampling schemes (see Subsection Implementation details). Our metrics to assess the performances are the average power and its standard deviation, as well as the average FDP and its standard deviation.

#### Stability study

To study the stability of the different variable selection methods with respect to perturbation in the input samples, we use ten times 10-fold cross-validation for a given simulated outcome **y**. At each step of the cross-validation we apply variable selection procedures to the training set, which contains 90% of the data. We extract the proportion of times each feature has been selected — referred to as the selection frequency — across the 10×10 iterations. We run the entire procedure with 10 different outcomes **y**, which gives a total of a 1000 different experiments.

#### Simulation difficulty evaluation

We classify the 100 iterations obtained with the LCD statistic in the statistics benchmark (see 3.4.1 for details) as easy, medium or hard post-hoc, based on the power obtained by the KO method for each outcome. To ensure a robust classification, we calculate the average power obtained for each simulation for a FDP ∈ [0, 0.3]. We then classify the simulations in 3 groups: hard (average power < 0.25), medium (average power ∈ 0.25, 0.7s) and easy (average power > 0.7). The thresholds were chosen arbitrarily.

#### Real data application

We apply KOPI with LSCIP and LCD methods for different target FDR levels (*q* ∈ {0.1, 0.2, 0.3, 0.5}) to the three real data sets presented in subsection Real data sets.

## Results

In the following, we present the results of the simulation study conducted to investigate the applicability of the KO framework to transcriptomic data.

### KO generation methods and statistics benchmark

We first compare the performance of different KO generation methods and statistics in the linear simulation setting.

We find that CI, LSCIP, MVR, and SDP have similar power, as illustrated in Fig. 1. Concerning the statistics, Fig. 1 clearly show better power for linear statistics (LCD, EN-CD, and MLR) — which was expected as we used a linear model to generate **y**. Over the 100 replicates, see supplementary figures S3 (d)), the EN-CD statistics have a lower power on average than the other two linear statistics for target FDR levels lower than 0.5. Our results also highlight that the EN-CD statistic is conservative with FDP lower on average than the target FDR level and the other three methods for *q* < 0.5 (see Fig. 1 (d)). On average, the four KO generation methods and statistics allow control of the number of false discoveries (see Fig. 1 (b-d)).

**Figure 1.**
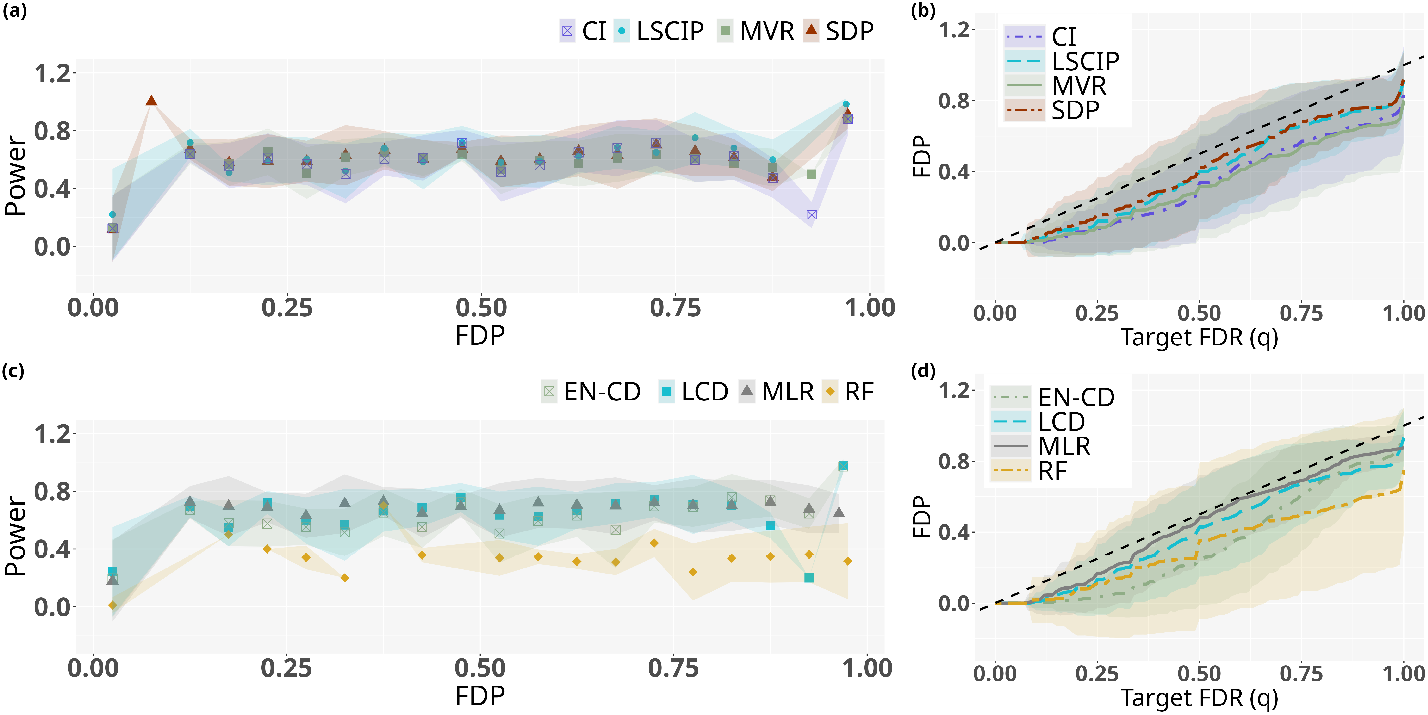
Comparison of the KO generation methods (a-b) and of different KO statistics (c-d). Outcomes are simulated from the CRUKPAP feature matrix using the linear simulation setting with *k* = 10 and *M* = 100 repetitions. (a-c) Power values are given as means over FDP bins of width 0.05. (b-d) False discovery proportion versus target false discovery rate. (a-b) Important variables are selected using the LCD statistics for all KO generation methods. CI: Conditional independence, LSCIP: Linear Sequential Conditional Independant Pairs algorithm, MVR: Minimum Variance-based reconstruction, SDP: semi-definite programming, see section Generation of KO features for details. (c-d) KO features are generated using the LSCIP method for all statistics. EN-CD: Elastic Net Coefficients Difference, LCD: Lasso Coefficients Difference, MLR: Maximum Likelihood Ratio, RF: Random Forest, see section KO test statistics for details.

We also compare the performance obtained with the different statistics in the interaction case, to account for the interacting nature of biological processes. All methods, even the RF nonlinear statistic, demonstrate a clear lack of performance (see Fig. 2 and Supp. Mat. C.1.3). We then conducted the same experiments on the AEGIS and BC datasets and found similar results (Supp. Mat. D), although we note that the performance of the KO methods are generally better on the BC cohort than on the others. Finally, it is notable that all our analyses, except for the BC data, exhibit large standard deviation intervals (see, for example, Supp. Fig. S3), indicating high variability in our results, which motivates the stability study below.

**Figure 2.**
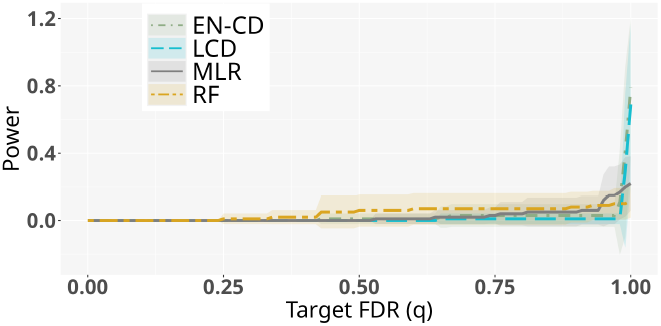
Comparison of different KO statistics in the interaction simulation setting. Outcomes are simulated from the CRUKPAP feature matrix with *k* =10 and *M =* 10 repetitions. Power values are given as means over FDP bins of width 0.05. EN-CD: Elastic Net Coefficients Difference, LCD: Lasso Coefficients Difference, MLR: Maximum Likelihood Ratio, RF: Random Forest, see section KO test statistics for details.

To conclude, this simulation study shows that all KO generation methods produce similar results, in contrast to other settings in which MVR or CI were shown to outperform SDP [20, 21]. Concerning the choice of statistics, MLR and LCD have shown the best performance in the linear simulation setting. In what follows, we will use the LSCIP method for KO generation because it does not make assumptions about the data structure and the LCD statistic since it is the most widely used in the literature.

After determining which KO generation method and which statistic to use, we compared the performance of the KO framework using the aggregation method KOPI (see subsection KO aggregation), and found that KOPI is as powerful as the standard KO procedure, and thus include KOPI in the following experiments (see Supp. Fig. S13 and Tab. 1).

**Table 1.**
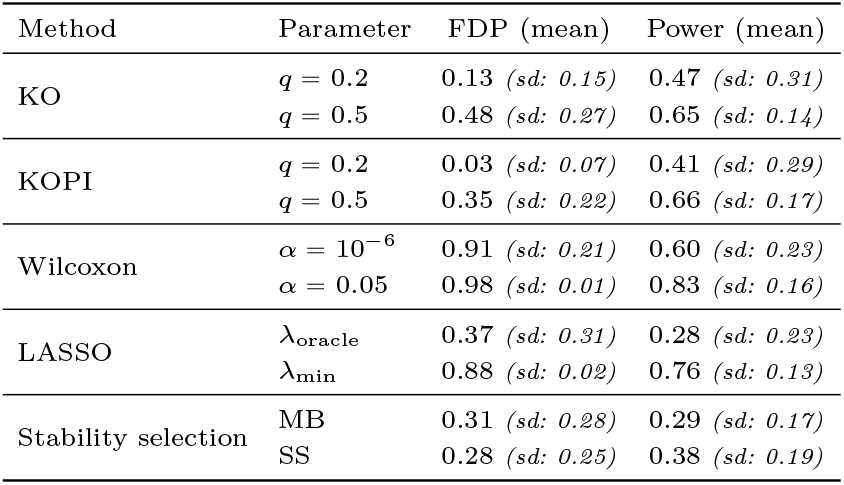
Comparison of the power and FDP obtained with different variable selection methods: the KO procedure and KOPI with *q* = 0.2 and *q* = 0.5, the LASSO with penalization parameters *λ*_min_ and *λ*_oracle_, the Wilcoxon rank-sum test with *α* = 10^−6^ and *α* = 0.05, and stability selection with the MB and SS sampling schemes. The mean and the standard deviation (sd) are computed for *M* = 100 linearly simulated **y** on the CRUKPAP features matrix.

### Comparison to baselines

We investigate how KO methods compare to other standard variable selection procedures. We consider the linear simulation setting and compare KO methods to a LASSO-penalized logistic regression, Wilcoxon rank-sum tests (which has shown a promising reduction in of the number of false discoveries compared to some other methods for large sample sizes in differential expression analysis [31, 35]), and stability selection.

Table 1 summarizes the power and FDP values obtained for the different methods (see also table S1 in the supplementary material for comparisons with additional methods). It underlines that the Wilcoxon rank-sum test and the LASSO computed with *λ*_min_ induce many false discoveries with an average FDP above 0.85. Of note, the wilcoxon test performs marginal tests while the lasso and KO framework consist in conditional testing. The high correlation between the features in our dataset makes the marginal testing of the wilcoxon test less accurate and might explain its very poor performances. The two KO-based methods demonstrate better power over FDP ratio than other methods. In particular, we optimize *a posteriori* the LASSO penalization parameter to maximize the power over FDP ratio (called *λ*_oracle_), and found that KO-based methods outperformed the LASSO even in the LASSO most ideal case. As in our previous benchmarks, standard deviations in Tab. 1 illustrate high variability of all methods.

To confirm our results, despite large standard deviation for both power and FDP, we compare the selection performance for larger sequences of false discoveries control parameters for the KO, KOPI and Wilcoxon rank-sum test and a sequence of penalization parameters for the LASSO. Regardless of the FDP interval considered after selection, the KO procedure and KOPI are more powerful than the LASSO and the Wilcoxon rank-sum test and show similar power (see Supp. Fig. S10 and Fig. S13). We also include Bayesian Variable Selection (BVS) for logistic regression [36] and a penalized Support Vector Machine (SVM) [37], which outperform both the Lasso and Wilcoxon test but remain less powerful than KO on average (see Supp. Mat. Sec. C.2.1).

Overall, these comparisons with baselines models demonstrate the ability of KO-based methods to improve selection power and to control the number of false discoveries.

### Stability study

Our previous results show that the KO framework controls the number of false discoveries while maintaining good power in the linear simulation setting. However, these performance metrics do not give any information about the impact of the stochasticity of the KO framework and the confidence we can have in the resulting selected subsets of genes. To fill this gap, we now turn to the study of feature selection stability that we use as a measure of reliability. Stability is defined as the extent to which two subsets selected by the same method resemble each other when the sample to which it is applied is disturbed [5]. To study variable selection stability, we simulate perturbation by subsampling of the initial samples (see Section Stability study for details) and analyze the selection frequency of each variable.

The results are given in Fig. 3. We find that LASSO-penalized logistic regressions, with penalization parameter chosen by cross-validation, selects too many variables and is unstable. In particular, out of a total of 749 × 10 genes, it has 2, 070 alternatively selected genes, *i.e*., genes that are neither always nor never selected–in other words, their selection frequency is not equal to 0 or 1. In addition, among the set of systematically selected genes, more than 40% are false discoveries.

**Figure 3.**
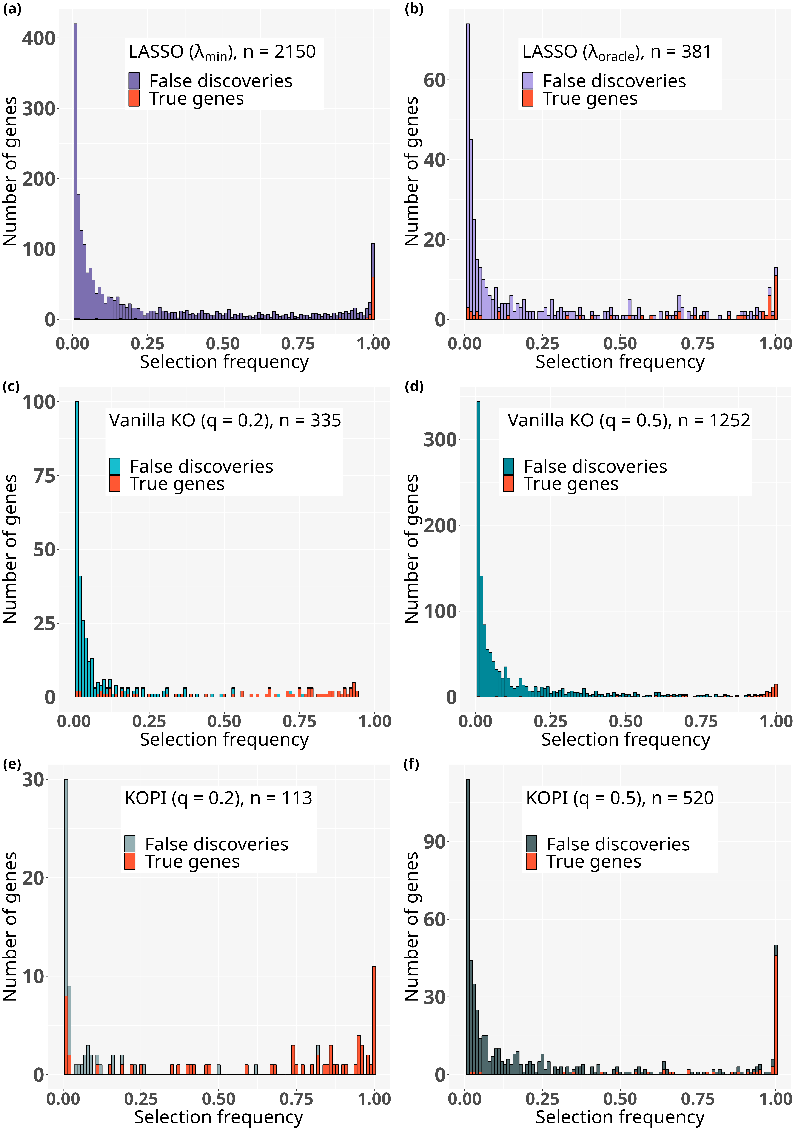
Histograms of the frequency of features selected *more than once* in different variable selection frameworks. Selection frequencies are computed over 10 iterations of ten-fold subsampling for each method and simulated outcome, in the linear simulation setting with *k* = 10 and the CRUKPAP features matrix. *n* gives the number of features among the 749 × 10 selected at least once across the 100 × 10 iterations. Features that were never selected across any iterations are not displayed. (a-b): Selection performed with LASSO-penalized logistic regressions, (a) for *λ* = *λ*_*min*_ or (b) *λ* = *λ*_*oracle*_ (c-d): Selection performed with KO for FDR level *q* = 0.2 (c) and *q* = 0.5 (d). (e-f): Selection performed with KOPI with target FDR level *q* = 0.2 (e) and *q* = 0.5 (f).

Conversely, KO-based methods tend to limit the selection of alternatively selected features, 335 for the vanilla KO and 102 for KOPI with *q* = 0.2. Taking *q* = 0.5 and thus relaxing the constraint on the number of false discoveries enables the selection of true causal features at almost each iteration but also leads to an increase of the number of alternatively selected features (1 237 with vanilla KO and 470 with KOPI). KOPI leads to fewer alternatively selected genes as the KO procedure, showing the ability of the aggregation to mitigate the effect of KO stochasticity while maintaining similar power. Yet, as illustrated in Figure S14 in supplementary material, KOPI is specifically designed to tackle KO stochasticity but remains sensitive to the sample (see (e-f) Fig. 3.

On the other hand, although stability selection methods most strongly limit the number of alternatively selected genes (see Supp. Fig. S15), they fail to identify many true genes.

Finally, the stability of the KOPI framework is equivalent to that of the oracle LASSO (set *a posteriori* to maximize the true-to-false discovery ratio), showing that KOPI is as good as the best possible selection set obtained with the lasso.

Table 2 summarizes the number of true and false discoveries found in Fig. 3. As expected, we find that the LASSO makes more true discoveries at the cost of a high number of false discoveries (proportion of true discoveries: 3.7%) while KOPI selects fewer features in order to control the FDR level. Nonetheless, KOPI discovers more true features than the stability selection methods. KOPI also demonstrates a higher proportion of true discoveries among the set of selected genes (for *q* = 0.2, 56.6% and for *q* = 0.5, 14.0%) than the LASSO (0.04%) and the oracle LASSO (13.7%). For the stability selection methods, the proportions (36.5% for MB and 44.4% for SS) are higher than those of KOPI with *q* = 0.5, but still lower than with *q* = 0.2.

**Table 2.** Number of true discoveries (TD) and false discoveries (FD) among systematically selected features (selection frequency = 1) and features selected at least once (freq > 0) for different variable selection methods. The selection frequency is computed over 10 iterations of ten-fold subsampling for each method and 10 simulated outcome **y** — resulting in 100 potential TD and 7390 FD. See Section 3.4.3 for experimental details. The outcomes **y** are simulated in the linear simulation setting with *k* = 10 and the CRUKPAP feature matrix.

| Method | Parameter | TD (FD)<br>(freq = 1) | TD (FD)<br>(freq > 0) |
| --- | --- | --- | --- |
| KO | $q = 0.2$ | 0 (0) | 74 (261) |
| | $q = 0.5$ | 15 (0) | 75 (1177) |
| KOPI | $q = 0.2$ | 11 (0) | 64 (49) |
| | $q = 0.5$ | 46 (4) | 73 (447) |
| LASSO | $\lambda_{\text{oracle}}$ | 11 (2) | 52 (329) |
| | $\lambda_{\text{min}}$ | 60 (48) | 80 (2070) |
| Stability selection | MB | 11 (3) | 27 (47) |
|  | SS | 15 (2) | 40 (50) |

Overall, our results illustrate both the conservative nature and the ability of KO-based methods to remove false discoveries specifically among systematically selected features. We also find that features selected by KOPI are more reliable than those selected by the KO framework or the LASSO, and that KOPI is more powerful than the stability selection methods.

### Influence of the correlation between features on performance

To study the causes of the differences in performance of the KO method between the different datasets (see Supp. Mat. D and Fig. 1), we compare the correlation matrices of the three cohorts (see Supp. Fig. S25) and find that the matrix data *X*_BC_ has a less correlated structure than both *X*_CRUKPAP_ and *X*_AEGIS_. This suggests that correlation between causal genes may reduce power.

To test this hypothesis, we introduce difficulty categories for the true causal genes (“easy”, “medium”, and “hard”, see Simulation difficulty evaluation for details), which depict how easy it is for the KO framework to find these 10 true causal variables. We categorize in Tab. 3 each of the 100 simulations used in section KO generation methods and statistics benchmark and in supplementary material D). Tab. 3 shows that the selection is easier with the breast cancer cohort (54 “easy” cases) than with the other two cohorts (0 and 7 “easy” cases for CRUKPAP and AEGIS cohorts).

**Table 3.**
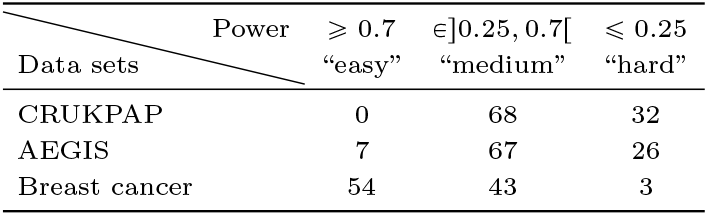
Classification of the performance of variable selection using the KO framework for different transcriptomic data sets. The outcomes **y** are simulated 100 times with, at each iteration, 10 randomly selected non null genes from a real feature matrix *X* (*X* ∈ {*X*_CRUKPAP_, *X*_AEGIS_, *X*_BC_} see Supp. Mat. D for details). The power is given as the average power obtained over an FDP interval: FDP ∈ [0, 0.3].

As an illustration, we compare in Fig. 4 the average power obtained over ten “easy” settings and ten “hard” settings for the CRUKPAP and the breast cancer cohorts. These curves point out that the CRUKPAP results demonstrate larger variability in performance than the breast cancer results. It also shows that the identity of the 10 non null features is the main source of performance variability. To quantify how correlation affects performance, we show in Fig. 5 the mean power as a function of the average pairwise correlation between the non null features. We find that the highest power values are reached for low pairwise correlations (< 0.2). It also depicts a general tendency of the KO framework to be more powerful on average for lower pairwise correlations. Ultimately, higher correlation in the data matrix makes selection harder in the KO framework. We have shown that selection performance and variability depend partly on the correlation between true causal features.

**Figure 4.**
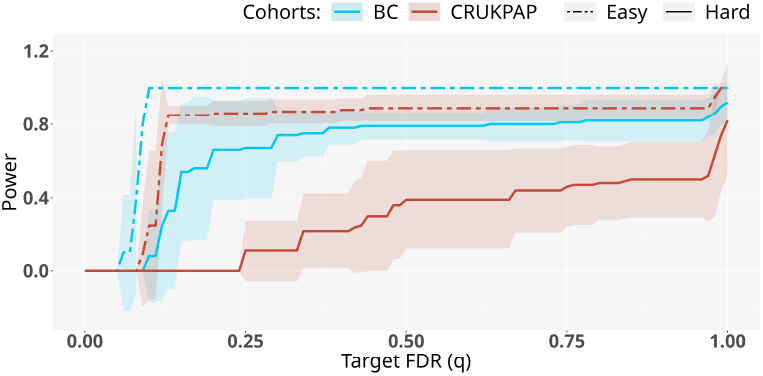
Illustration of the performance variation of the KO procedure (LSCIP + LCD) The power is obtained for the 10 most difficult (continuous line) or the 10 easiest case (dotted line) out of *M* = 100 different outcomes simulations using the linear simulation setting with *k* = 10 and from the CRUKPAP (red) or the breast cancer (blue) feature matrix.

**Figure 5.**
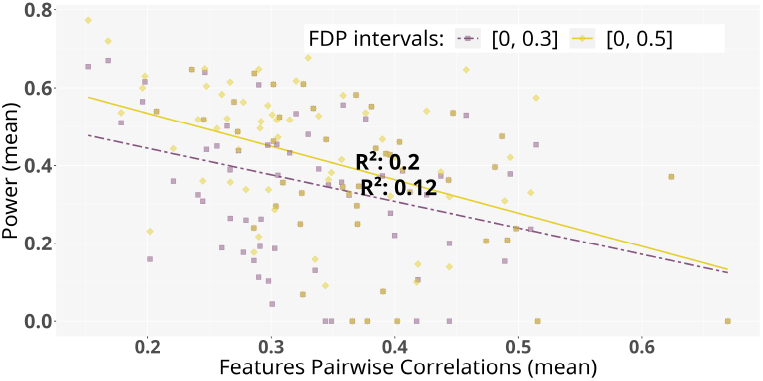
KO variable selection power depending on the pairwise correlation between the 10 real important features. Simulations are conducted with the linear simulation setting on the CRUKPAP feature matrix. The mean power is computed for two different FDP intervals: [0, 0.3] (purple dots) and [0, 0.5] (yellow dots). Each dot corresponds to a different simulation experiments. Continuous and dotted lines represent the linear regression fit of the mean power versus pairwise correlation for the two different FDP intervals.

### Application to real data

To evaluate the applicability of KO-based procedures in real settings, we apply KOPI to perform variable selection with real transcriptomic data from CRUKPAP, AEGIS and the breast cancer data sets, but this time **y** is set to real outcomes— for which we thus do not know the true causal genes. For CRUKPAP and AEGIS, we aim to select variables involved in cancer diagnosis (cancer/no cancer). In the breast cancer study, we are looking for genes that can discriminate breast cancer biomarkers: ER status (positive/negative), PgR status (positive/negative), and HER2 status (amplified/normal).

We find that for lung cancer predictions, ER and PgR status predictions, no genes are selected with *q* = {0.1, 0.2, 0.3}. For *q* set to 0.5, only 3 genes are selected in the CRUKPAP cohort and 7 in the AEGIS cohort (see Supp. Tab. S2). For the prediction of ER status in the breast cancer cohort, only one gene, the ESR1 gene (coding for the estrogen receptor) is selected. Likewise, in the sets of genes selected as involved in HER2 status (6 genes for *q* = 0.3 and 13 genes for *q* = 0.5), we retrieve the gene coding for the HER2 protein (ERBB2). The fact that no genes are selected for the PgR status prediction is consistent with the results of the original breast cancer study where the PgR status was shown to be the hardest to predict from PGR expression levels [29].

Finally, we also note that the ERBB2 gene is less correlated to other genes as compared to ESR1 and PGR (see Supp. Fig. S26 and Fig. S27). This lower level of correlation might in part explain why the KO-framework selects more genes in the HER2 prediction case, in good agreement with our results of section Influence of the correlation between features on performance. Overall, these results illustrate the conservative nature of KOPI in real settings.

## Discussion

In this paper, we studied the applicability of the KO framework to variable selection problems on transcriptomic data. We focused on a common use case where the goal is to find genes associated with a phenotype of interest—such as a disease. This is in contrast to most applications of KO which are conducted in a regression setting. We benchmark different statistics and KO generation methods, and found that all KO generation methods yield similar results in our simulations. MLR and LCD statistics showed the best performance in the linear simulation setting while being, like other statistics, powerless in the interaction simulation setting. Our extensive simulation study shows that KO succeed in significantly reducing the false discovery rate compared to standard feature selection procedures. While using KO can also reduce statistical power, the KO framework exhibits an improved power over false discovery ratio as compared to standard approaches in our simulations. Finally, we also show that using KO aggregation strategies allows to mitigate the instability of gene selection, a problem often disregarded when comparing feature selection procedures, but nevertheless of major importance. We note however that those results were obtained on a relatively simple simulation framework, which might not fully reflect the complexity of real biological scenarios.

Applying KO to real transcriptomic data also demonstrates the limitations of the framework. First, the KO framework tends to be very conservative, leading in several of the real examples we studied to very few or sometimes no single discovery. However, these real life examples are taken from previous research where the authors select tens or hundreds of genes to construct clinical classifiers that achieve good performance, suggesting that there is signal in the data and features to be selected. While highlighting the conservative nature of the KO procedure, this discrepancy also indicates that the genes selected in the original studies potentially contain many false discoveries.

Second, we show that while KO performs well when features are moderately correlated, its power decreases when the correlation between the causal features reaches very high values. While KO are constructed precisely to perform variable selection for correlated features, our analyses show this is effective only up to a point. Interestingly, we find that the level of correlation between the true causal features impacts the performance of the KO selection procedure more strongly than the correlation between causal features and non-causal features.

Third, we focused mostly on the case of *k* = 10 true causal features in our simulations. Qualitatively, we obtain similar results with *k* = 30 (*i.e*. KO still outperforms other feature selection methods, see Supp. Fig. S17). However, we note that the performance of the KO methods deteriorates when the number of causal features increases, see Supp. Fig. S16. Similarly, the performance of the KO methods also declines as the number of non-causal features increases (see Supp. Fig. S18). Nevertheless, even with all available features, KO still outperforms the LASSO method (see Supp. Fig. S19). How the KO framework behaves for larger values of the number of samples *n* remains to be investigated. An additional difficulty of working in high dimensions lies in the computational cost of the generation of the KO matrix, which increases rapidly as the dataset size increases [18].

Finally we find that variable selection performance collapses when the link between gene expression and outcome is not linear. It is however very likely that for many biological processes, the dependency between features and the outcome is not linear. This difficulty is not specific to KO, as all of the classical methods we compare have very low performances in our simulations (see Supp. Fig. S11). Our results thus highlight the need for novel methodological developments to tackle feature selection for correlated data in nonlinear classification settings. To tackle this challenge, one strategy could be to group genes that belong to similar biological pathway, as already implemented successfully many times [38]. To do so, one strategy could be to use kernel based method in combination with KO (see [39] for a similar approach in a univariate setting). Another strategy could be to use deep learning methods that offer a flexible framework and might help in non-linear context. Although our first attempts have given very poor results — probably due to the small sample size of our cohorts (see Fig. S6 and S9) — approaches using reinforcement learning strategies might allow to overcome sample size challenges.

Another interesting result of our study is to underline the instability of classical gene selection procedures, as previously observed [5]. While standard KO do not improve selection stability—notably because of the inherent stochasticity of the KO framework—we find that using KO aggregation methods such as KOPI [14] greatly improves the stability of the selection procedure (Fig. 3), while maintaining similar statistical power. We note however that KOPI calibration is not ideal and the FDR control appears very conservative (see Tab. 2), such that very few discoveries are obtained with a target FDR rate of 0.2. In our simulations, we obtain a low number of false discoveries (< 5%) at the cost of limited power (41%) for a target rate of 0.2. Yet, these results demonstrate a better power-to-FDP ratio than other approaches.

Overall, our study further demonstrates the difficulty of classification tasks with transcriptomic data. We nevertheless show that KO can constitute an interesting strategy, in particular to improve gene selection stability. Our study shows how KO can be applied to transcriptomic dataset, leading the way to more sophisticated approaches such as combining KO to biological pathway analyses.

## Keypoints

- Knockoffs-based methods demonstrate a better power-to-false discovery proportion ratio than other commonly used variable selection methods in the context of high-dimensional classification from transcriptomic data.
- The stability of the selection is improved by using an aggregation scheme that mitigates the effects of knockoffs stochasticity while maintaining the same power.
- Both real-world applications and simulations highlight several limitations of the knockoff framework: its conservative nature, its sensitivity to the correlation structure and the number of true causal features, and its poor performance in nonlinear classification settings.

## Author contributions statement

A.F., C.A.A and F.M. conceived the experiment(s), J.C. and J.L. conducted the experiment(s), and J.C., A.F., C.A.A and F.M. analysed the results. J.C. wrote the manuscript, A.F., C.A.A and F.M. reviewed the manuscript.

## Authors

Chloe Agathe Azencott is a professor, Florian Massip is a permanent researcher, Julie Cartier is a PhD student and Johanna Lagoas is a master intern. All work at mines paris PSL. Adeline Fermanian is a researcher at Califrais.

## Data Availability

The code for all experiments is available in the following Github repository (https://github.com/Julie-cartier/RNAKnockoffs). The breast cancer gene expression data are available from the NCBI Gene Expression Omnibus under accession GSE96058. The AEGIS gene expression data are also available in the NCBI Gene Expression Omnibus under accession number GSE80796.

## Acknowledgments

This work was carried out with the financial support of Carnot M.I.N.E.S, as part of its leading project CARINGS project, dedicated to health engineering. This work was also supported by the French Agence Nationale de la Recherche (ANR-19-P3IA-0001). The authors would like to thank the Cancer Research UK, Royal Papworth Hospital and the co-authors of De Biase et al [27] for access to the CRUKPAP cohort data (EGA project number: EGAD50000000333).

