## Supplementary Material for "Statistical knockoffs improve biomarker discovery from transcriptomic data"

**F Supplementary Material: Application to real data** **24**

### A Supplementary Material: The Knockoff framework

#### A.1 Overview of the KO framework

In the paper, KO refers to the versatile Model-X Knockoffs [1] proposed to generalize the original framework [2] to high dimension [1]. All the definitions, properties and results of this section have been introduced in [1] to build the KO framework. For the sake of completeness, they are briefly recalled here.

Let $\mathbf{y} \in \{0,1\}^n$, $\mathbf{y} = (y_1, \dots y_n)^T$ be the random variables representing the outcomes and $X = (X_{.,1}, \dots, X_{.,p})$ be the set of features such that for any $j \in \{1, \dots p\}$, $X_{.,j} = (X_{1,j}, \dots, X_{n,j})^T \in \mathbb{R}^n$. For the sake of clarity, in the following $X_{.,j}$ will be noted $X_j$. We assume that the data pairs $(X_{i,.}, Y_i)_{i \in \{1,\dots,n\}}$ are i.i.d. Let $\mathcal{H}_0$ denote the index of null features such that for all $j \in \mathcal{H}_0, \mathbf{y} \perp\!\!\!\perp X_j | X_{-j}$, where $X_{-j}$ is the matrix $X$ without the $j^{\text{th}}$ column and $\perp\!\!\!\perp$ denotes independence between variables. The goal of variable selection is to identify the set of non null features, *i.e.* the set $\mathcal{H}_0^c = \{1, \dots, p\} \backslash \mathcal{H}_0$.

In addition to recovering the set of non-zero features, the KO framework ensures that the false discovery rate (FDR) is controlled. Therefore, running the KO framework results in the selection of a set of variables $\widehat{\mathcal{S}} \subset \{1, \dots, p\}$ such that:

$$\text{FDR} = \mathbb{E}\left[\frac{|\widehat{\mathcal{S}} \cap \mathcal{H}_0|}{|\widehat{\mathcal{S}}| \vee 1}\right] \leqslant q, \tag{1}$$

where $q$ is the target FDR level.

The KO framework relies on the construction of false copies of every feature in the data matrix $X$ that will be used as controlled variables. These copies are then gathered to form the KO matrix $\widetilde{X} = (\widetilde{X}_1, \dots, \widetilde{X}_p) \in \mathbb{R}^{n \times p}$.

**Definition 1** (Model-X knockoffs)**.** $\widetilde{X} = (\widetilde{X}_1, \dots, \widetilde{X}_p)$ *is the KO matrix of the original features* $X$ *if* $\widetilde{X}$ *satisfies the following two properties:*

1. *For any subset* $S \subset \{1, ..., p\}$,
$$(X, \widetilde{X})_{\text{swap}(S)} \overset{d}{=} (X, \widetilde{X})$$
*where* $(X, \widetilde{X})_{\text{swap}(S)}$ *is obtained by swapping* $\widetilde{X}_l$ *and* $X_l$ *in* $(X, \widetilde{X})$ *for any* $l \in S$, *and* $\overset{d}{=}$ *denotes equality in distribution.*

2. *Conditionally on* $X$, *the Model-X knockoff is independent of the features, that is,* $\widetilde{X} \perp\!\!\!\perp \mathbf{y}|X$.

These new features are used to compute statistics for every features of the data matrix. Let $W_j = w_j((X, \widetilde{X}), \mathbf{y})$ denote the statistic of the $j^{\text{th}}$ feature for some function $w_j$. The function is chosen so that a high value of the statistic provides evidence that the corresponding feature is non-zero. In addition, for any $j \in \{1, \dots, p\}$, $W_j$ must respect the flip-sign property, given below.

**Definition 2** (flip sign property)**.** *A statistic* $W_j = w_j((X, \widetilde{X})$ *satisfies the flip sign property if for any subset* $S \subset \mathcal{H}_0$ *and for any* $j \in \{1, \dots, p\}$*:*

$$w_j((X, \widetilde{X})_{\text{swap}(S)}, \mathbf{y}) = \begin{cases} -w_j((X, \widetilde{X}), \mathbf{y}), \textit{ if } j \in S \\ w_j((X, \widetilde{X}), \mathbf{y}) \textit{ otherwise.} \end{cases}$$

Finally, Candès et al. [1] have demonstrated that, under these assumptions, $\widehat{\mathcal{S}} = \{j : W_j \geqslant \tau_+\}$, where

$$\tau_+ := \min \left\{ t > 0 : \frac{1 + |\{j : W_j \leqslant -t\}|}{|\{j : W_j \geqslant t\}|} \leqslant q \right\}$$

for a chosen $q$, allows FDR control (1).

In order for the KO features to be used as powerful control features in the computation of statistics, the KO framework requires KO features to be as distinguishable as possible from the original features. Here, the power quantifies the ability of variable selection methods to retrieve non-null features and is defined by:

$$\text{Power} = \frac{|\widehat{\mathcal{S}} \cap \mathcal{H}_0^c|}{|\mathcal{H}_0^c|}.$$

The task is, then, to build KO matrices and statistics that satisfy the above properties (swap Eq. 1 in Definition 1, conditional independence Eq. 2 in Definition 1 and flip-sign 2) to enable control of the number of false discoveries, that achieve good power.

#### A.2 Generation of KOs features

To construct a KO matrix $\widetilde{X}$ that satisfies the conditional independance property (Eq. 2 in Definition 1), it is sufficient to never use the outcome vector $\mathbf{y}$ in the construction. In contrast, building a KO matrix that has the same correlation structure as the original feature matrix (Eq. 1 in Definition 1) while remaining distinguishable from the original matrix is challenging. Various methods have been proposed to address this problem, we present some of them below.

**Gaussian methods** The most common methods assume that the feature matrix $X$ follows a Gaussian distribution. Under this assumption, it is possible to compute a parametrized family of distributions of $\widetilde{X}|X$, from which one can sample a valid KO for the original features [1]. Several methods to select a parameter that achieve a good power have been proposed.

In the family of Mean Absolute Correlation (MAC) Gaussian methods, the KO features are computed to minimize the correlation between a feature $X_j$ and its knockoff $\widetilde{X}_j$ [1]. The different MAC-approaches refer to the type of optimization problem and/or the type of constraints used to compute the KO matrix: the semidefinite program (SDP), the equicorrelated program, and the approximate semidefinite program (ASDP) [1]. These methods have shown empirical robustness to deviation from the Gaussian assumption.

The MAC-based KO frameworks suffer from a tendency to identify some KO of non-null original features as important features in the model instead of the non-null features themselves, which is known as the "reconstructability effect" [3]. To tackle this, a new family of Gaussian KOs generation methods called Minimum Reconstructability (MRC) has been developed. A first approach is to maximize the conditional variance $\mathbb{V}ar(X_j|X_{-j}, \widetilde{X})$, for any $j \in \{1, ..., p\}$, which is known as the minimum variance-based reconstructability method (MVR). An alternative is to maximize the entropy of $(X, \widetilde{X})$ to have the least statistical dependence between X and Y, called maximum entropy method (ME) [4].

To improve power, another line of work explores the construction of KO that satisfy conditional independence between a feature $X_j$ and its KO $\widetilde{X}_j$, given the other covariates $X_{-j}$ — referred to as conditional independent (CI) KO. Their existence was initially demonstrated only for very specific structures of the covariance matrix $\Sigma$, like tree graphical models or sparse graphs [5]. The framework was then extended to less specific covariance matrices [6].

**SCIP methods** The Sequential Conditional Independant Pairs algorithm proposed in [1] allows for the construction of valid KOs with no further assumption on the distribution of the features $X$. In practice, a linear empirical version of this algorithm is used, called the Linear Sequential Conditional Independant Pairs algorithm (LSCIP) [7, 8].

**Algorithm 1** LSCIP Pseudo-Algorithm

```
for j = 1 to p do
    Fit a LASSO model on (X_{-j}) with X_j as outcomes
    Compute residuals ε = (ε_1, ..., ε_p), ε = X_j − X̂_j where X̂_j is the predicted value of X_j with the regression model
    Permute the residuals vectors randomly. Denote by ρ_p the permutation of {1, ..., p}
    Compute X̃_j = X̂_j + ε_{ρ_p(j)}
end for
return X̃
```

Different versions of this algorithm exist, for instance in [9], the authors use principal component analysis before applying a regression model.

In addition, methods based on deep learning have been proposed to generate KO features [10, 11], which we do not consider in this article since we have few samples in our data sets.

The previous list of methods to compute valid knockoffs is not exhaustive. Having prior knowledge about the data, such as the specific distribution of the covariates [12, 13] can allow the development of more specific and adapted methods. However, to our knowledge, no methods specifically tailored for transcriptomic data have been developed.

#### A.3 KO test statistics

We have seen that the KO framework ensures that we do not make too many false discoveries. However, we also need to find the maximum number of variables truly correlated with the output, that is, have a good power. Ensuring that the KO matrix is properly constructed with the correct properties is not sufficient to guarantee a good power. It also requires methods to compute statistics that efficiently discriminate important from non important features.

Most KO statistics suitable for high-dimensional data and classification frameworks are actually equal to differences between variable importance metrics. More precisely, the idea consists in applying a machine learning model to $(X, \widetilde{X})$ with target $\mathbf{y}$, choose a variable importance metric associated to the model, and compute the difference between the importance measure assigned to a feature and that assigned to its KO.

For linear settings, an intuitive option is to use the absolute value of the LASSO-penalized logistic regression coefficients (LASSO Coefficient Difference (LCD)) [2, 1] as the variable importance. An other option proposed in [2] is to use the largest penalty at which a feature enters the LASSO model (Lambda Difference (LD)). Given that elastic-net regularization has been shown to provide more stable selection with highly correlated data [14], and that transcriptomic data are correlated, we also consider using elastic-net penalized logistic regression to compute statistics (EN-CD).

Likewise, variable importance in nonlinear models, such as tree-based models, can be used to define statistics. For example, the default random forest based statistic (RF statistic) in the *knockoff* [3] package is computed as a the difference between Gini impurity importance [15]. In [16], the importance metric is given by quantifying at each step of the construction of the boosted tree the contributions of each feature to risk reduction (risk reduction in boosting (RRB)). Other boosted tree-based statistics are explored in [9] where variable importance is derived from different metrics such as frequency (the number of times a feature is used as a splitting variable), gain (the gain in accuracy given to a feature), or SHAP value (combining frequency and gain).

In [13], the authors propose a generic strategy to compute variable importance relying on measuring accuracy drop when a variable is replaced by a fake version, computed by some permutation or linear combination with its KO counterpart. There are two standard ways to create these fake variables called swap and swap integral, which can be applied with any classification model, but is implemented in *knockpy* [3] for random forests and deep learning models.

The DeepPINK (Deep feature selection using paired input nonlinear knockoff) [17] method combine a multi-layer perceptron model and an additional pairwise coupling layer. The importance metric is given by the weight of the coupling layer. To improve power, they used weighted variable importance as a statistic where the weights are computed throughout the layers of the MLP.

The literature also contains statistics that can not directly be expressed as variable difference. The LASSO Signed Max (LSM)[2] is an example of such methods. The LSM statistic gives the maximum between the largest penalty parameter at which a feature and its KO counterparts enter a LASSO penalized logistic regression model which sign depends on which one enters the model first. An other different approach is the masked likelihood ratio (MLR) [18], based on the estimation of log-likelihood ratios, which accounts for high correlation between a feature and its KO by lowering the absolute value of their statistic when the correlation is large.

All of the described statistics can be applied to biomarker discovery in transcriptomic data and do not require any prior knowledge.

##### A.4 KO aggregation

The stochastic nature of KO construction methods means that applying the same KO construction method twice to the same data matrix will generate different KO matrices. This leads to different subsets of selected variables at each iteration, as illustrated in Fig. S17. Addressing this issue is precisely the purpose of KO aggregation. The idea is to apply the KO procedure multiple times and to aggregate "knowledge" from these multiple runs to select variables. There exist several methods of aggregation, for example frequency [19] and quantile aggregation [20].

In particular, [7] propose to use $\pi$- statistics for each variables — considered as p-values in [20] under the assumptions that statistics are independent. The $\pi$-statistic of a feature $j$ is defined by:

$$\pi_j = \begin{cases} \frac{1+|k:W_k \leqslant -W_j|}{p}, \text{ if } W_j > 0 \\ 1, \text{ otherwise} \end{cases}$$

and gives evidence against the null hypothesis. The authors leverage the idea of controlling the joint error rate of these statistics to propose a probabilistic control of the FDP at level $\alpha$ — $\mathbb{P}(\text{FDP} \leqslant q) \geqslant 1 - \alpha$, where $q$ is the user-required bound of FDP. They demonstrate that this result holds for aggregated $\pi$-statistics. The resulting KOPI (Knockoff-$\pi$) framework [7] has shown promising performance compared to previous aggregation methods even for $n < p$ in both experimental and genomic real applications.

### B Supplementary Material: Material and methods

#### B.1 Outcomes simulations

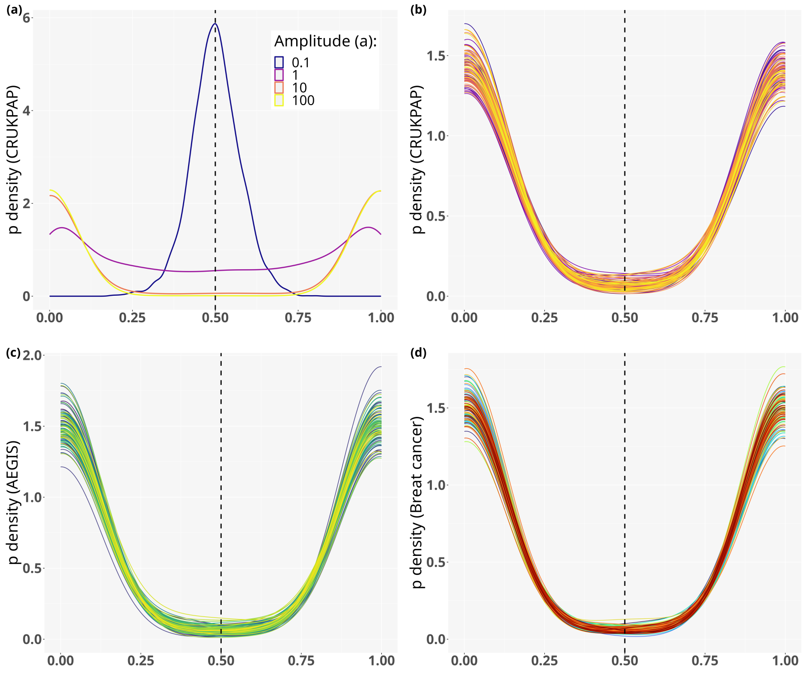

Figure S1: Density of the values of $p \in \mathbb{R}^n$, where $p = \sigma(X_{\text{data}}\boldsymbol{\beta}) = \frac{1}{1+e^{-X_{\text{data}}\boldsymbol{\beta}}}$ obtained for different magnitudes $a$ in (a) and for the different real datasets in (b), (c), (d). (a): The density is computed for each $a$ with 10 simulated $p$. Our goal is to see if a threshold of 0.5 divides properly the two distinct classes $y = 0$ and $y = 1$. We conclude that choosing $a = 10$ allow both a good and balanced separation of classes. (b), (c), (d): The density is computed for 100 different values of $\beta$ with, respectively, $X_{\text{CRUKPAP}}$, $X_{\text{AEGIS}}$, and $X_{\text{BC}}$.

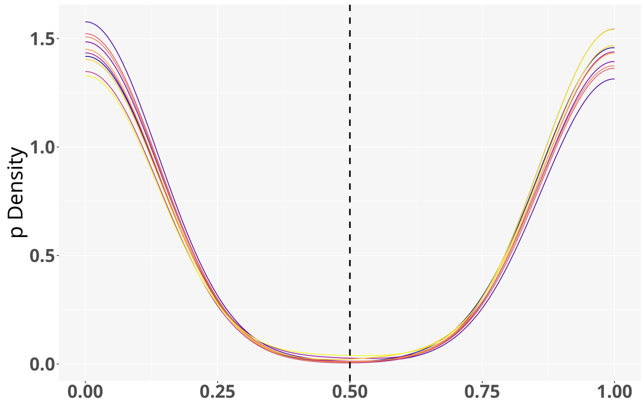

Figure S2: Density of the values of $p \in \mathbb{R}^n$ in the interaction simulation setting, where $p_i = \left(1 + e^{-f(X_{\text{CRUKPAP}_{i,\cdot}}, \boldsymbol{\beta})}\right)^{-1}$ and $f(X_{i,\cdot}, \boldsymbol{\beta}) = \sum_{j=1}^{\frac{p}{2}} \boldsymbol{\beta}_j X_{i,j} X_{i,2j}$ where $\beta_j$ is the $j^{\text{th}}$ coordinate of $\boldsymbol{\beta}$. The density values are obtained for $a = 100$, for the 10 simulated $\boldsymbol{\beta}$ corresponding to the vectors used in the interaction experiments (see Section Outcome simulation for details).

##### B.2 Simulation details

###### B.2.1 KO generation

We use the Python package *knockpy* (v1.3.0) [3] to implement Gaussian-based knockoffs (MVR, ME, CI, SDP). For the LSCIP algorithm, we use the *glmnet* (v4.1-7) R package [21]. At each iteration, the penalization parameter $\lambda$ is chosen to minimize the mean cross-validated error which is by default computed with the mean squared error over 10-fold cross-validation.

###### B.2.2 Knockoff statistics

The LASSO-based statistics (LD (lambda-difference), LSM and LCD) are computed using the R package *knockoff* [1] (v0.3.6). For the LCD statistic, the penalization parameter $\lambda$ is obtained by minimizing the mean cross-validated error over 10 folds, where the error is the deviance—difference of log-likelihoods. To compute both the LD and LSM statistics, 500 different values of $\lambda$ are used. We use the *glmnet* package to compute the EN-CD statistics. The parameter $\lambda$ is computed over 10-fold cross-validation to minimize the mean cross-validated error where by default the error is given by the deviance metric in logistic regression; the second penalization parameter $\alpha$ is set to 0.5. The MLR statistic is implemented in *knockpy.*

The swap and swap integral tree based statistics are computed using *knockpy.* The GINI-based random forest statistics are implemented in *knockoff.* To compute RRB statistics, we use the code provided by [16]. It relies on the use of the R package *mboost* [22] (v.2.9-7). We use trees as base learners, 200 for the parameter *max.mstop* which gives the maximum number of boosting iterations, and set to 10 the number of cross-validation folds used to get the optimal number of iterations. In *mboost* the *cvrisk* function, used to perform cross-validation, minimizes the empirical risk, which is by default the weighted sum of the negative log-likelihood loss function. Finally, the MLP based statistics are computed with the *knockpy* package.

For all use of the vanilla KO framework the parameter $\tau_+$ is computed with the *knockoff* package.

###### B.2.3 KOPI : Knockoff aggregation scheme

KOPI relies on two important parameters: the expected number of false discoveries $q$ and the confidence level parameter $\alpha$. Unless otherwise specified, for all the experiments with KO aggregation, $\alpha$ is set to 0.1. Note that changing the internal parameters of the KOPI method produced similar results both on real and simulated data, see Supp. Fig. S12. We use the code provided by [7] to perform KO aggregation. We keep the same algorithm parameter $k_{\max} = \lfloor \frac{p}{50} \rfloor$ and the harmonic mean as aggregation function. We set $B = 1\,000$. In our experiments, the number of iterations used to perform aggregation is $D = 100$

###### B.2.4 Baselines simulations

We apply the Wilcoxon rank-sum test on already preprocessed data as in [23] with the R *stats* package (v.4.2.0). We correct for multiple hypotheses testing with the Benjamini-Hochberg (BH) [24] procedure for different target FDR levels. We use the *glmnet* package to run LASSO-penalized logistic regression. The parameter $\lambda_{\text{min}}$ corresponds to the penalization obtained by minimizing the mean cross-validated error over 10 folds, where the error is the deviance. The parameter $\lambda_{\text{oracle}}$ is chosen *a posteriori* and corresponds to the penalization term that maximizes the ratio between true discoveries and false discoveries in a grid of values ranging from 0 to 1 in increments of 0.001. We use the same parameters $\lambda_{\text{oracle}}$ and $\lambda_{\text{min}}$ to run LASSO in the stability study. Stability selection with LASSO-penalized logistic regression is implemented with the R package *stabs* (v0.6-4). Two sampling schemes were considered: the original method of Meinhausen and Bühlmann [25] (MB) and the complementary pairs approach proposed by Shah and Samworth [26] (SS). For both schemes, we set the cutoff parameter to 0.75 and the upper bound of the per-family error rate at 1. The number of subsamples is set to 100, corresponding to 50 complementary pairs for the SS sampling scheme. LASSO-penalized logistic regression models are fit using *glmnet*.

**Additional methods for variable selection performance comparison** To perform Bayesian Variable Selection (BVS), we use the R *varbvs* package (v.6.10). We set the family argument to "binomial" and use the default setting for all other arguments. The penalized Support Vector Machine (SVM) is computed with the R *hdsvm* package (v.1.0.2)[27]. We use the Smoothly Clipped Absolute Deviation (SCAD) penalty. For the penalization parameter "lambda", we use a sequence of 100 values between 1 and $10^{-4}$. We set the quadratic penalty term to 0.01.

### C Supplementary Material: Additional experiments on the CRUK-PAP cohort

#### C.1 KO generation methods and statistics comparison

##### C.1.1 Additional Figures: Statistics and KO generation methods comparison

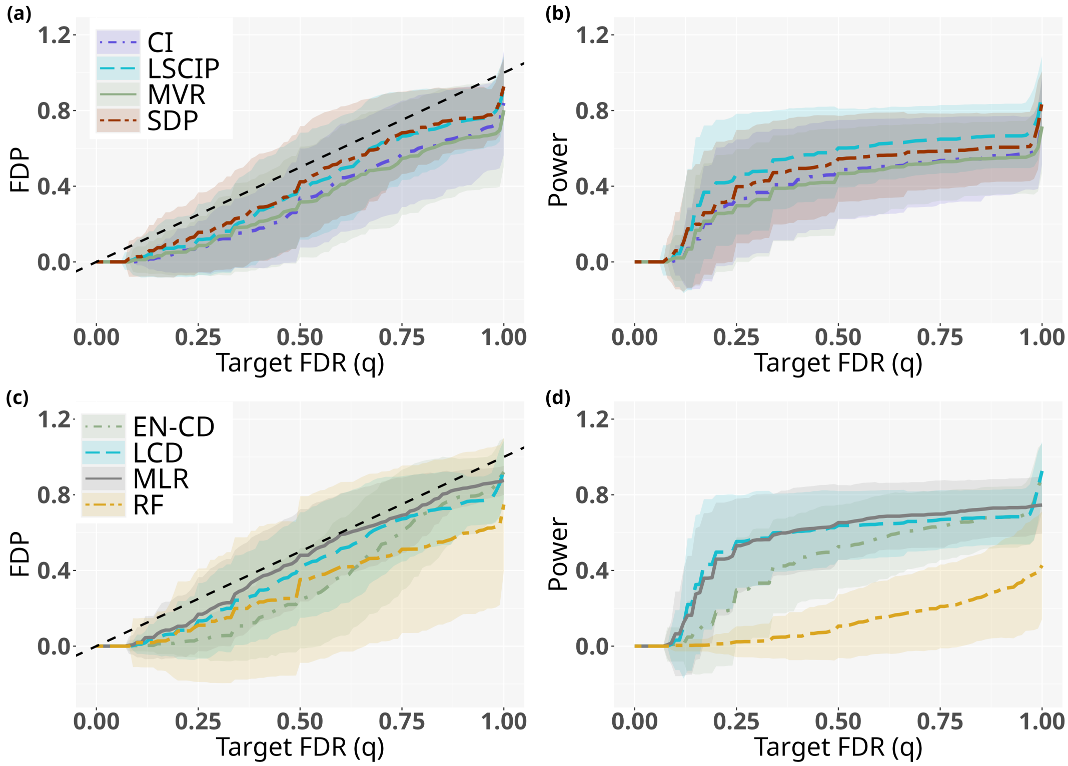

Figure S3: **Comparison of the KO construction methods and statistics.** Outcomes are simulated from the CRUKPAP feature matrix using the linear simulation setting with $k = 10$ and $M = 100$ repetitions. (a-c) False discovery proportion versus target false discovery rate. (b-d) Power versus target false discovery rate. (a-b) Important variables are selected using the LCD statistics for all KO generation methods. CI: Conditional independence, LSCIP: Linear Sequential Conditional Independant Pairs algorithm, MVR: Minimum Variance-based reconstruction, SDP: semi-definite programming, see section A.2 for details. (c-d) KO features are generated using the LSCIP method for all statistics. EN-CD: Elastic Net Coefficients Difference, LCD: Lasso Coefficients Difference, MLR: Maximum Likelihood Ratio, RF: Random Forest, see A.3 for details.

##### C.1.2 Figures: Preliminary comparisons of different statistics in the linear simulation setting

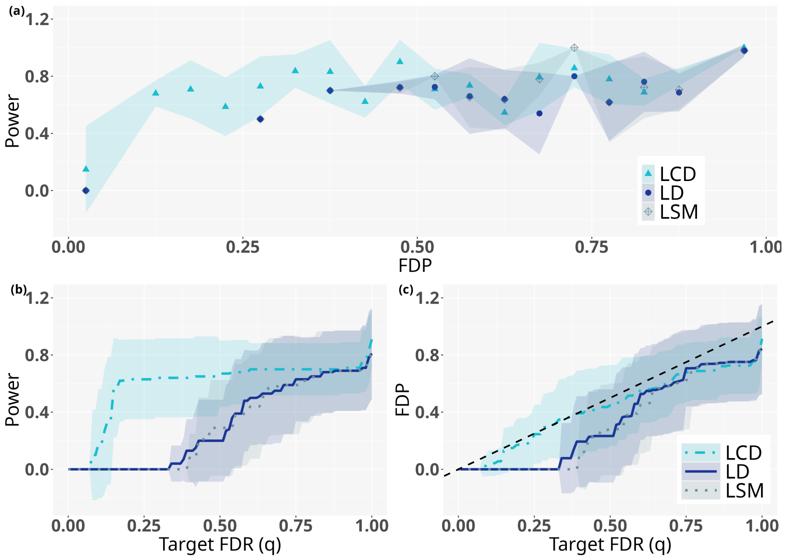

Figure S4: **Comparison of different LASSO logistic regression based statistics** Outcomes are simulated from the CRUKPAP feature matrix using the linear simulation setting with $k = 10$ and $M = 10$ repetitions. KO features are generated using the LSCIP algorithm. (a) Power versus false discovery proportion. Power values are given as means over FDP bins of width 0.05. (b) Power versus target false discovery rate. (c) False discovery proportion versus target false discovery rate. LCD: Lasso Coefficients Difference, LSM: Lasso Signed Max: Maximum Likelihood Ratio, LD: Lambda Difference, see A.3 for details.

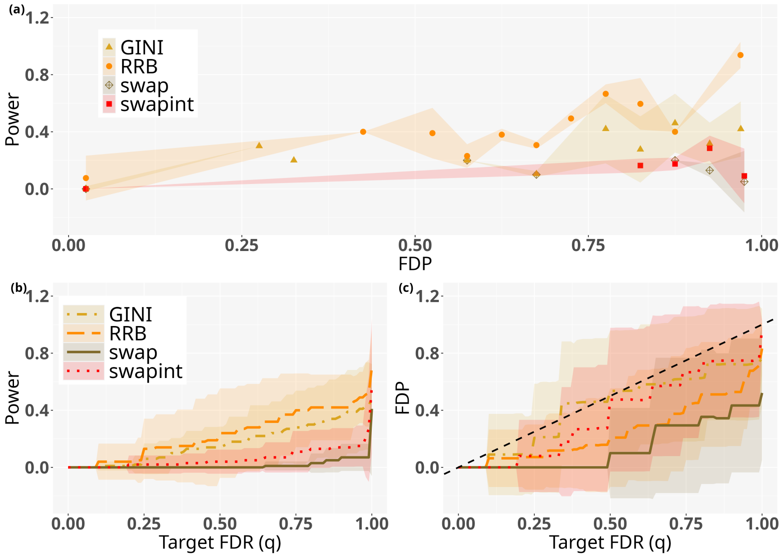

Figure S5: **Comparison of different decision tree-based statistics.** Outcomes are simulated from the CRUKPAP feature matrix using the linear simulation setting with $k = 10$ and $M = 10$ repetitions. KO features are generated using the LSCIP algorithm. (a) Power versus false discovery proportion. Power values are given as means over FDP bins of width 0.05. (b) Power versus target false discovery rate. (c) False discovery proportion versus target false discovery rate. Gini Importance, RF: Random Forest (swap/swap integral), RRB: Risk Reduction in Boosting see A.3 for details.

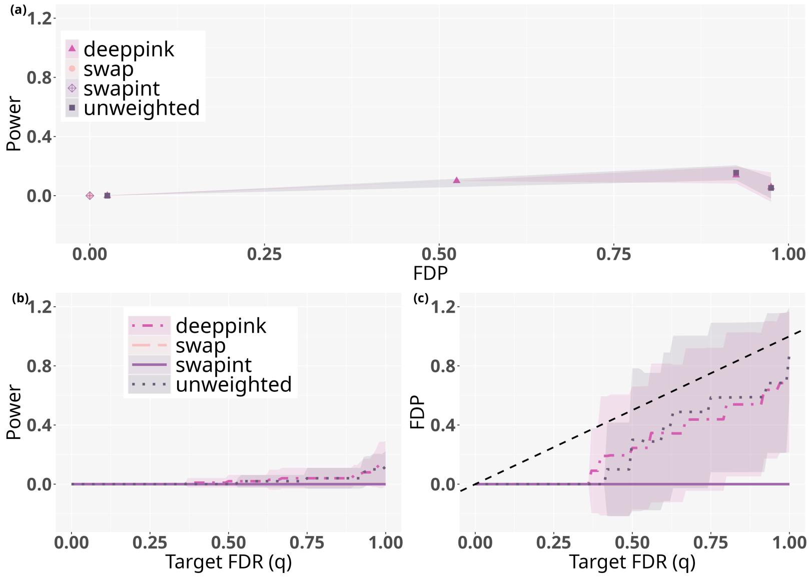

Figure S6: **Comparison of different MLP-based statistics.** Outcomes are simulated from the CRUKPAP feature matrix using the linear simulation setting with $k = 10$ and $M = 10$ repetitions. KO features are generated using the LSCIP algorithm. (a) Power versus false discovery proportion. Power values are given as means over FDP bins of width 0.05. (b) Power versus target false discovery rate. (c) False discovery proportion versus target false discovery rate. DeepPINK, MLP: Multi-Layer Perceptron (swap/swap integral), unweighted DeepPINK, see A.3 for details.

##### C.1.3 Figures: Additional results in the interaction simulation setting

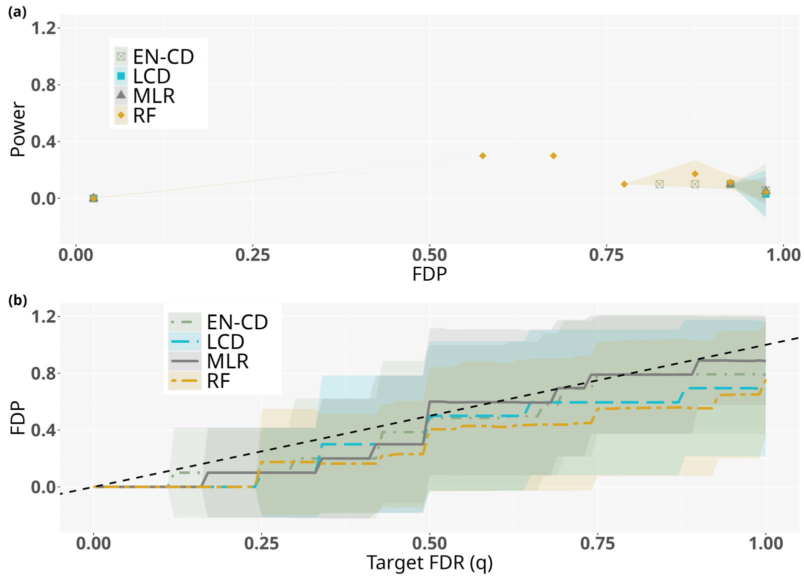

Figure S7: **Comparison of different KO statistics in the interaction simulation setting.** Outcomes are simulated from the CRUKPAP feature matrix using the interaction simulation setting with $k = 10$ and $M = 10$ repetitions. KO features are generated using the LSCIP algorithm. (a) Power versus false discovery proportion. Power values are given as means over FDP bins of width 0.05. (b) False discovery proportion versus target false discovery rate. EN-CD: Elastic Net Coefficients Difference, LCD: Lasso Coefficients Difference, MLR: Maximum Likelihood Ratio, RF: Random Forest, see A.3 for details.

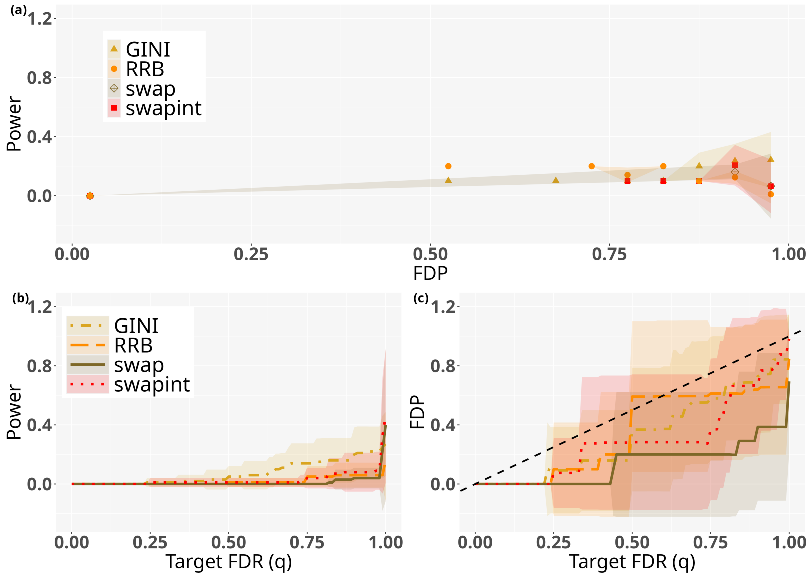

Figure S8: **Comparison of different decision tree-based statistics in the interaction simulation setting.** Outcomes are simulated from the CRUKPAP feature matrix using the interaction simulation setting with $k = 10$ and $M = 10$ repetitions. KO features are generated using the LSCIP algorithm. (a) Power versus false discovery proportion. Power values are given as means over FDP bins of width 0.05. (b) Power versus target false discovery rate. (c) False discovery proportion versus target false discovery rate. Gini Importance, RF: Random Forest (swap/swap integral), RRB: Risk Reduction in Boosting see A.3 for details.

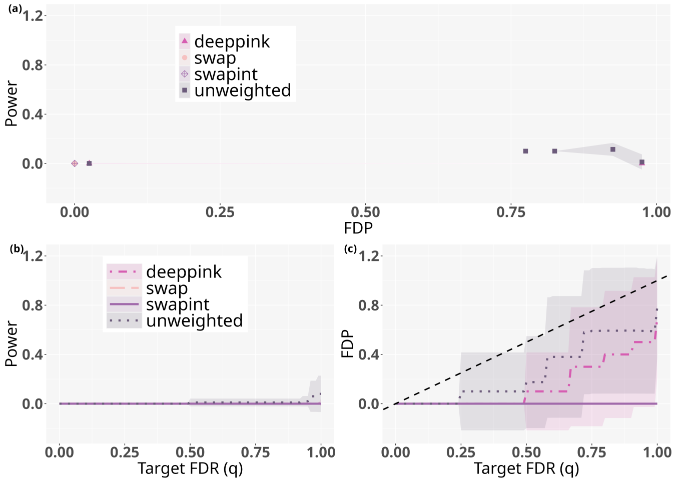

Figure S9: **Comparison of different deep learning-based statistics in the interaction simulation setting.** Outcomes are simulated from the CRUKPAP feature matrix using the interaction simulation setting with $k = 10$ and $M = 10$ repetitions. KO features are generated using the LSCIP algorithm. (a) Power versus false discovery proportion. Power values are given as means over FDP bins of width 0.05. (b) Power versus target false discovery rate. (c) False discovery proportion versus target false discovery rate. DeepPINK, MLP: Multi-Layer Perceptron (swap/swap integral), unweighted DeepPINK, see A.3 for details.

#### C.2 Comparison to other methods

##### C.2.1 Comparison in the linear simulation setting (KO framework, LASSO, Wilcoxon rank-sum test, Bayesian variable selection and penalized SVM )

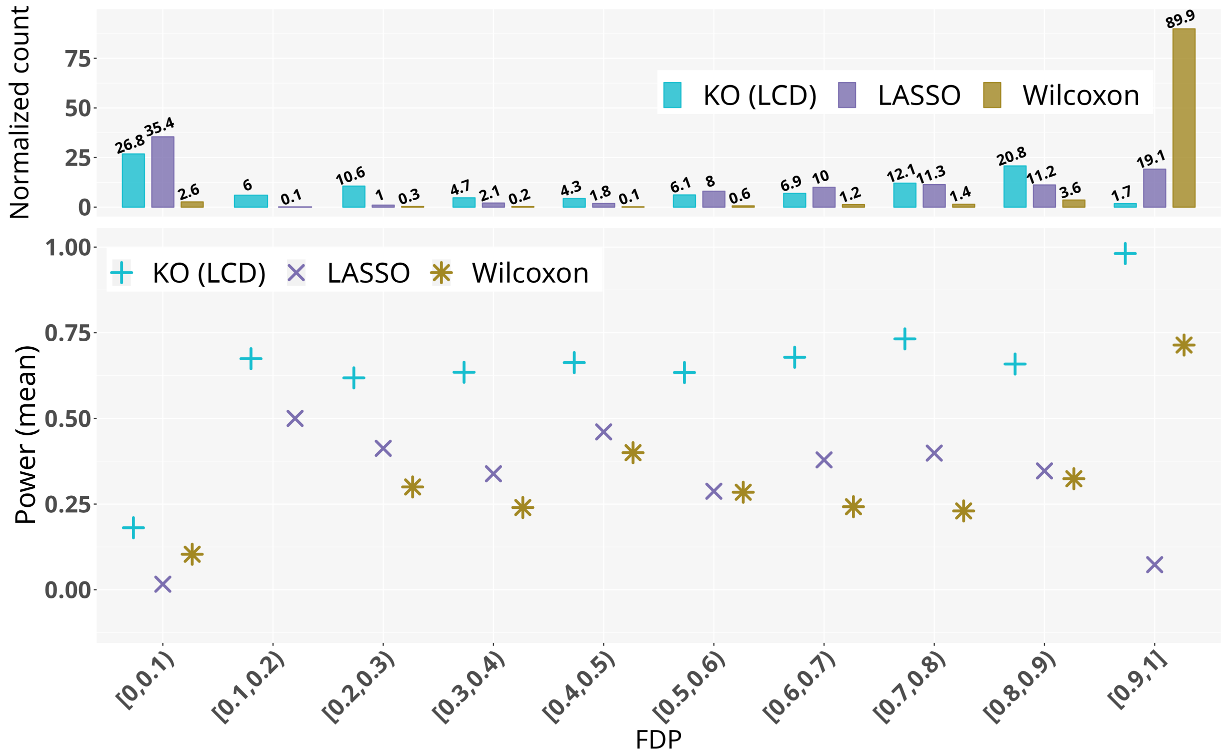

Figure S10: **Comparison of different feature selection methods: the KO procedure, the lasso penalized logistic regression, and the Wilcoxon rank-sum test** Outcomes are simulated from the CRUKPAP feature matrix using the linear simulation setting with $k = 10$ and $M = 100$ repetitions. The normalized count gives the number of points in the corresponding FDP interval –*i.e.* the number of points on which the mean power was calculated– normalized by the length of the sequence of parameters for each method (KO: $q$, LASSO: $\lambda$, Wilcoxon: $\alpha$) (for details see Subsection Experimental setups).

**Additional methods: Bayesian Variable Selection and penalized SVM** We use the same experimental setting as described in Section 3.4.2. From the Bayesian Variable Section in logistic regression, we obtain for each feature a posterior inclusion probability — that is the probability of each feature to be included in the model. We consider two approaches to perform selection. In the first one, we take the 10 variables with the highest posterior inclusion probabilities. For the second approach, we sequentially add features in the set of selected features based on their probability of inclusion — in decreasing order —. We vary the number of selected features from 1 to 749. The "oracle" set of selected features is the one that maximizes the ratio between true discoveries and false discoveries.

For the SCAD penalized SVM, the penalization parameter $\lambda_{SCAD_{\min}}$ is chosen by cross validation — 5-fold — to minimize the mean cross-validated error.

| Method | Parameter | FDP (mean) | Power (mean) |
| --- | --- | --- | --- |
| KO | $q = 0.2$ | 0.13 *(sd: 0.15)* | 0.47 *(sd: 0.31)* |
| | $q = 0.5$ | 0.48 *(sd: 0.27)* | 0.65 *(sd: 0.14)* |
| KOPI | $q = 0.2$ | 0.03 *(sd: 0.07)* | 0.41 *(sd: 0.29)* |
| | $q = 0.5$ | 0.35 *(sd: 0.22)* | 0.66 *(sd: 0.17)* |
| BVS | 10 | 0.46 *(sd: 0.21)* | 0.49 *(sd: 0.19)* |
| | oracle | 0.20 *(sd: 0.19)* | 0.43 *(sd: 0.21)* |
| SVM | $\lambda_{SCAD_{\min}}$ | 0.29 *(sd: 0.27)* | 0.47 *(sd: 0.31)* |

Table S1: Comparison of the power and FDP obtained with different variable selection methods: the Bayesian Variable Selection (BVS), the KO procedure and KOPI with $q = 0.2$ and $q = 0.5$, the SVM with penalization parameters $\lambda_{SCAD_{\min}}$. The mean and the standard deviation (sd) are computed for $M = 100$ linearly simulated $\mathbf{y}$ on the CRUKPAP features matrix.

##### C.2.2 Comparison in the linear interaction setting (KO framework, LASSO, and Wilcoxon rank-sum test)

We use the same experimental setup as in the linear simulation setting (see Section Comparison between KO, LASSO, Wilcoxon rank-sum test, KOPI and Stability Selection for details)

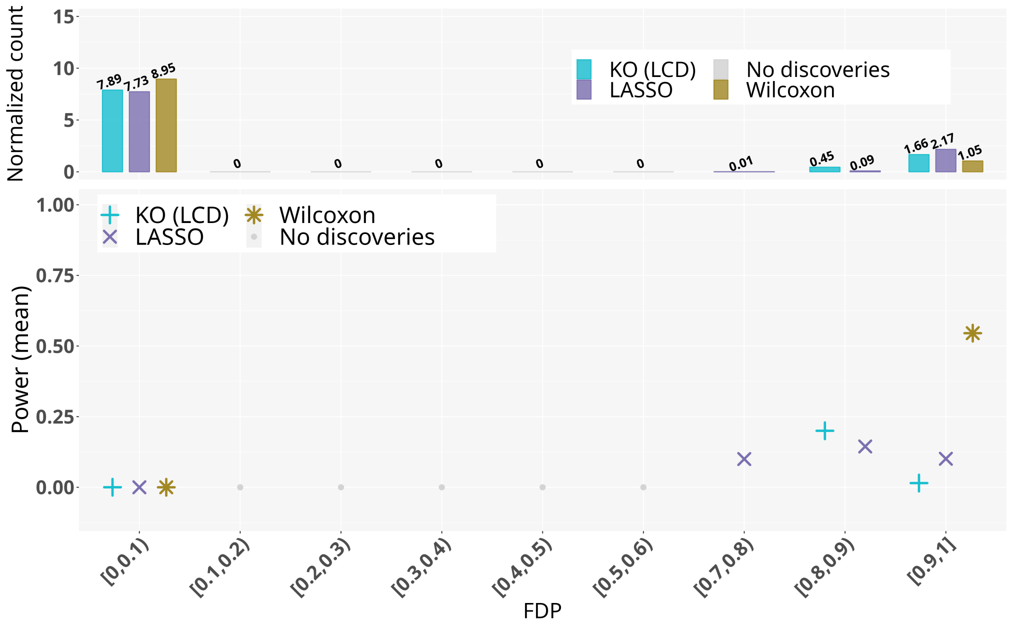

Figure S11: **Comparison of different feature selection methods: the KO procedure, the lasso penalized logistic regression, and the Wilcoxon rank-sum test** Outcomes are simulated from the CRUKPAP feature matrix using the interaction simulation setting with $k = 10$ and $M = 100$ repetitions. The normalized count gives the number of points in the corresponding FDP interval –*i.e.* the number of points on which the mean power was calculated– normalized by the length of the sequence of parameters for each method (KO: $q$, LASSO: $\lambda$, Wilcoxon: $\alpha$) (for details see Subsection Experimental setups).

#### C.3 Additional Figures: KOPI study

##### C.3.1 Performance of KOPI for different FDP control levels

We use the same simulation settings described in Section B.2.3 in the supplementary material, except that we vary the $\alpha$ parameter.

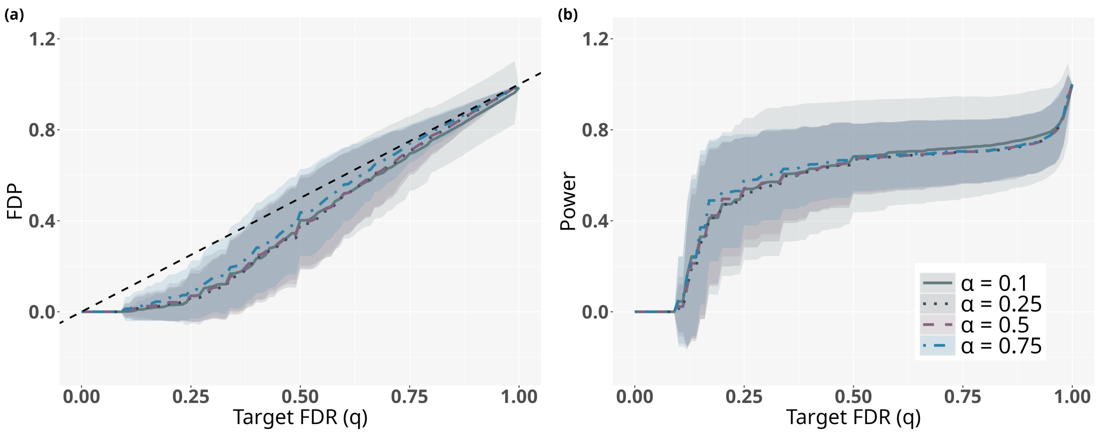

Figure S12: **Comparison KOPI performance for different FDP control levels** $\alpha$**.** Outcomes are simulated from the CRUKPAP feature matrix using the linear simulation setting with $k = 10$ and $M = 100$ repetitions. (a) False discovery proportion versus target false discovery rate. (b) Power versus target false discovery rate. KOPI is computed with $\alpha \in \{0.1, 0.25, 0.5, 0.75\}$.

##### C.3.2 Performance comparison between the KO framework and KOPI

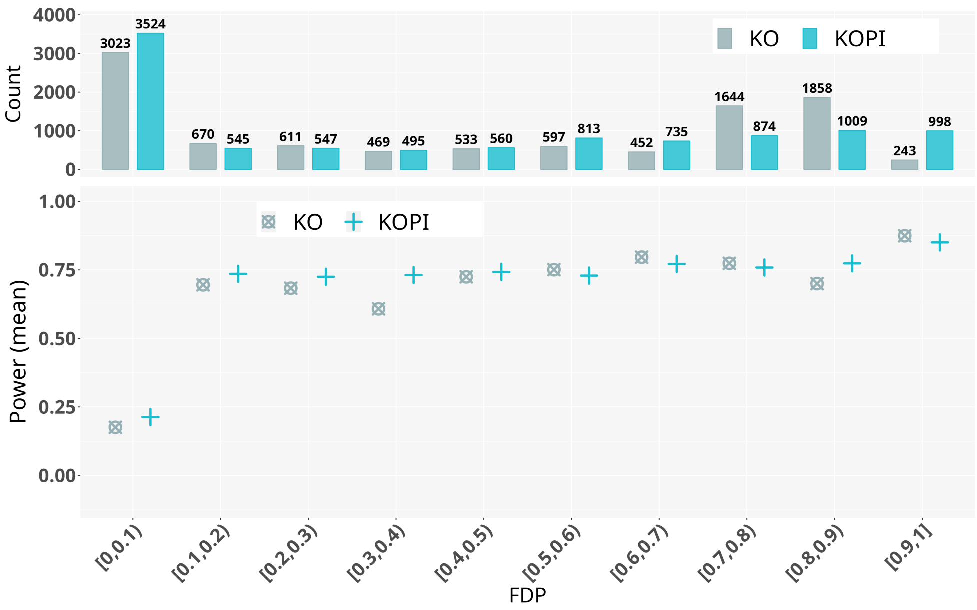

Figure S13: **Comparison of different feature selection methods: the KO procedure and KOPI** Outcomes are simulated from the CRUKPAP feature matrix using the linear simulation setting with $k = 10$ and $M = 100$ repetitions. The count gives the number of points in the corresponding FDP interval—*i.e.* the number of points on which the mean power was computed.

##### C.3.3 KOPI effect on KO stochasticity

To assess KOPI impact on KO stochasticity, we simulate one outcome $y$ in the linear simulation setting. We apply KOPI and the KO framework 100 times for $q \in \{0.2, 0.5\}$. We then compute the selection frequency of each variable.

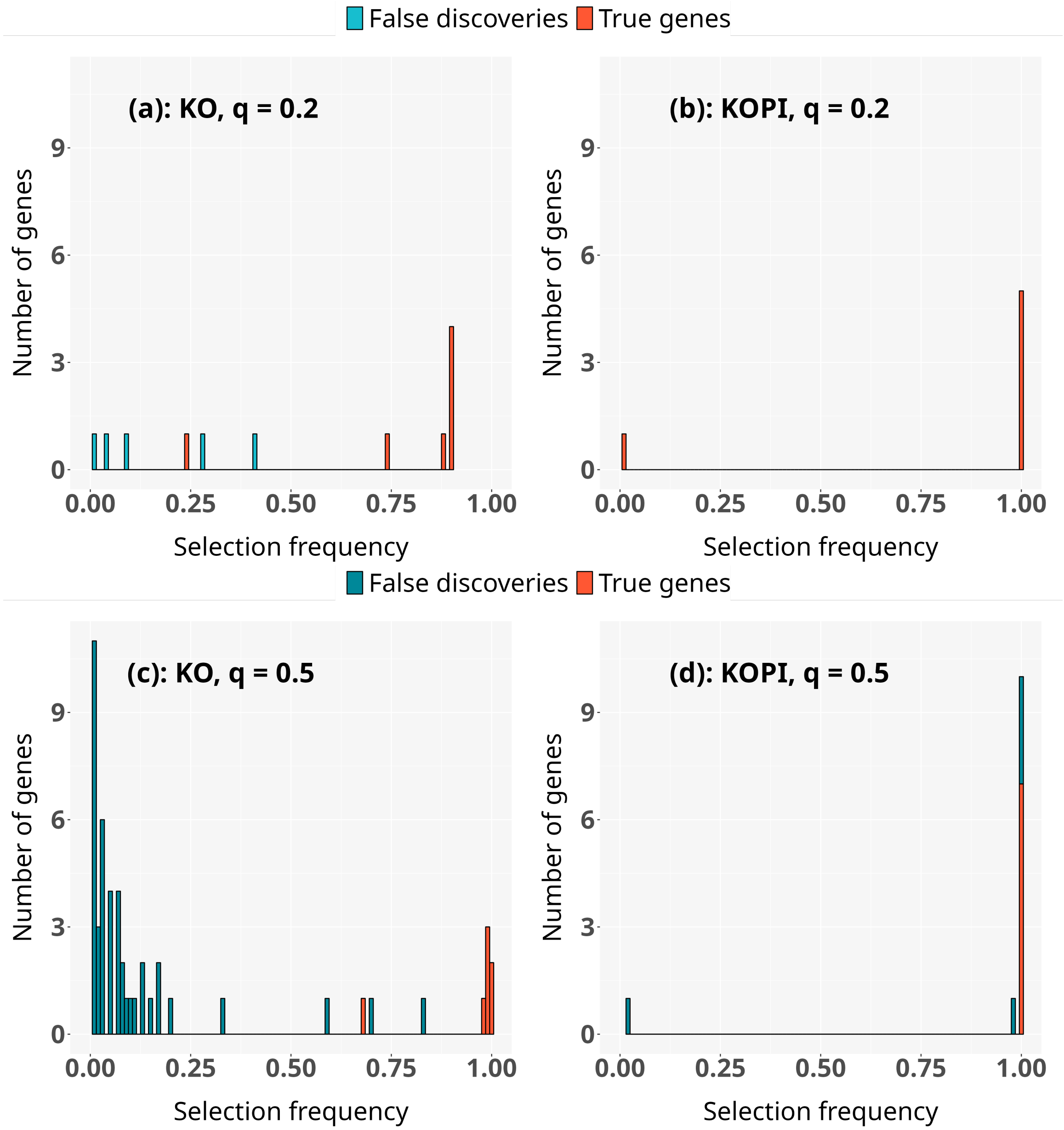

Figure S14: **Histograms of the frequency of features selected *more than once* with KOPI and the KO framework**. Selection frequencies are computed over 100 iterations for one simulated outcome, in the linear simulation setting with $k = 10$ and the CRUKPAP features matrix. Features that were never selected across any iterations are not displayed. (a-c): Selection performed with KO for FDR level $q = 0.2$ (a) and $q = 0.5$ (c). (b-d): Selection performed with KOPI with target FDR level $q = 0.2$ (b) and $q = 0.5$ (d).

#### C.4 Additional Figures: Stability study of the stability selection methods

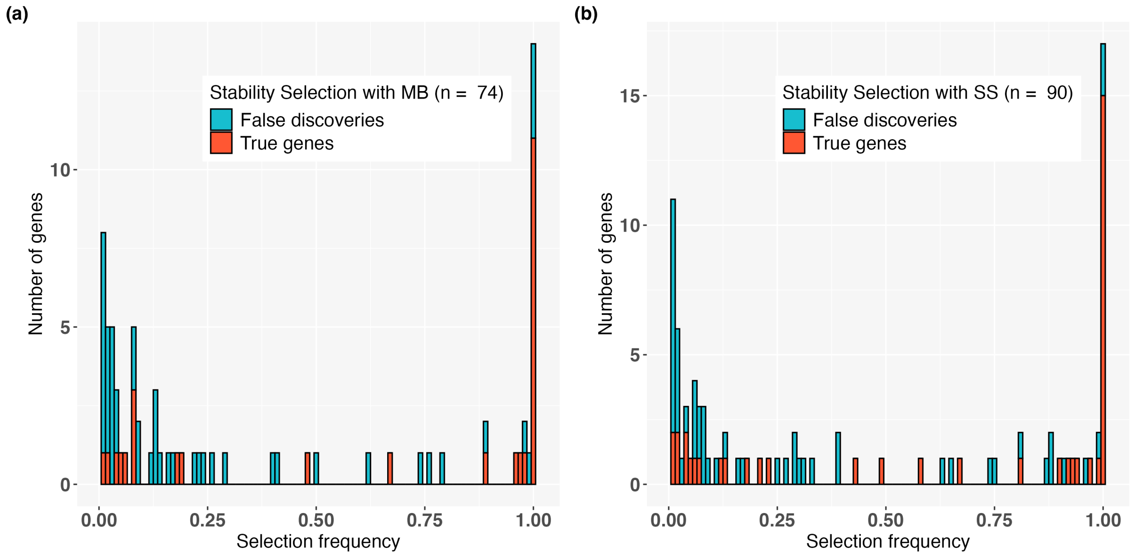

Figure S15: **Histograms of the frequency of features selected *more than once* with two stability selection methods**. Selection frequencies are computed over 10 iterations of ten-fold subsampling for each method and simulated outcome, in the linear simulation setting with $k = 10$ and the CRUKPAP features matrix. $n$ gives the number of features among the $749 \times 10$ selected at least once across the $100 \times 10$ iterations. Features that were never selected across any iterations are not displayed. (a-b): Stability selection performed with the MB (a) and SS (b) sampling schemes.

#### C.5 Effect of the number of non null features on the performance

We use the same experimental setup as in the linear simulation setting (see Section Comparison between KO, LASSO, Wilcoxon rank-sum test, KOPI and Stability Selection for details) for the comparison between different methods.

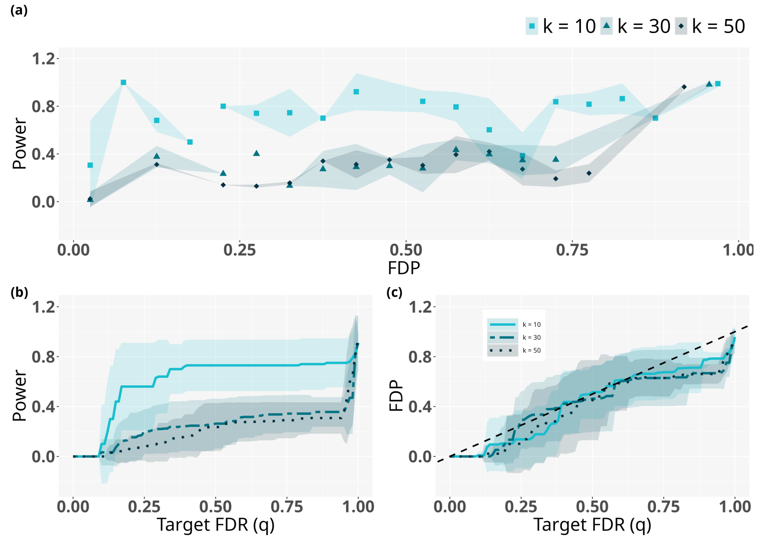

Figure S16: **Comparison of the KO procedure performance for different numbers of non-null features.** Outcomes are simulated from the CRUKPAP feature matrix using the linear simulation setting with $k \in \{10, 30, 50\}$ and $M = 10$ repetitions. (a) Power versus false discovery proportion. Power values are given as means over FDP bins of width 0.05. (b) Power versus target false discovery rate. (c) False discovery proportion versus target false discovery rate.

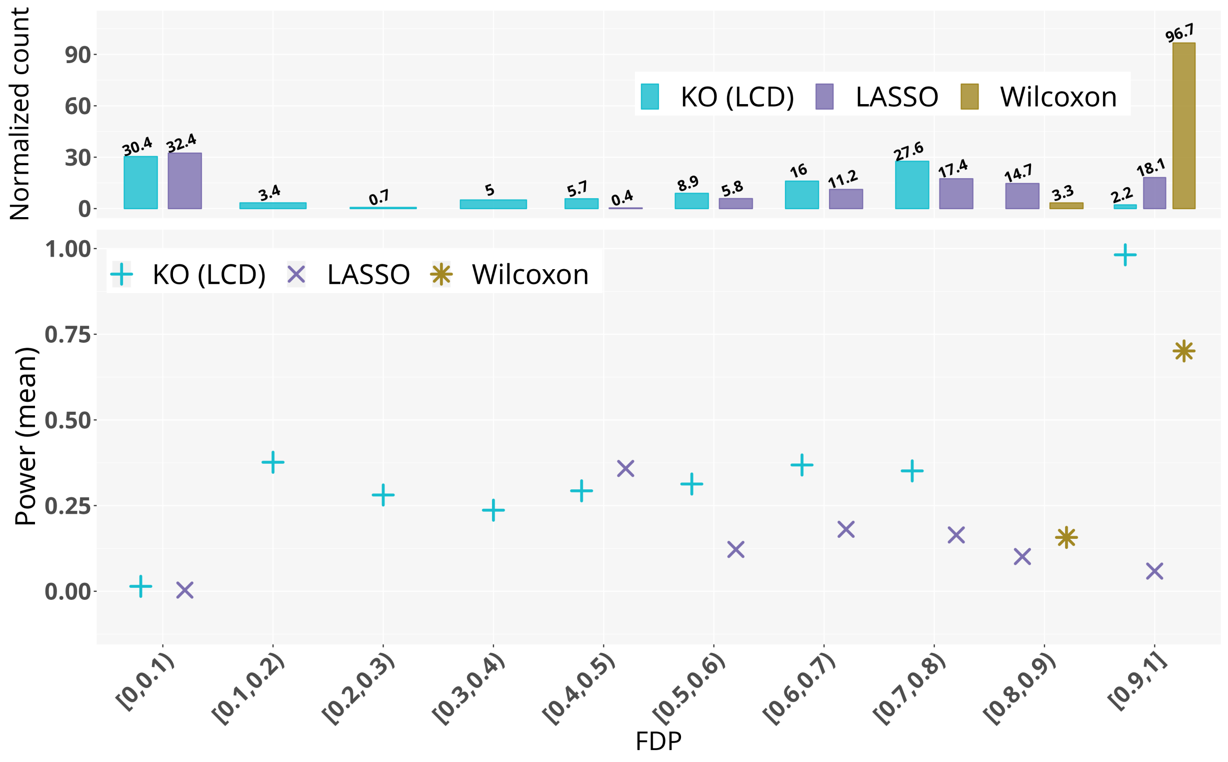

Figure S17: **Comparison of different feature selection methods: the KO procedure, the lasso penalized logistic regression, and the Wilcoxon rank-sum test** Outcomes are simulated from the CRUKPAP feature matrix using the linear simulation setting with $k = 30$ and $M = 100$ repetitions. The normalized count gives the number of points in the corresponding FDP interval—*i.e.* the number of points on which the mean power was calculated—normalized by the length of the sequence of parameters for each method (KO: $q$, LASSO: $\lambda$, Wilcoxon: $\alpha$) (for details see Subsection Experimental setups).

#### C.6 Effect of the number of null features on the performance

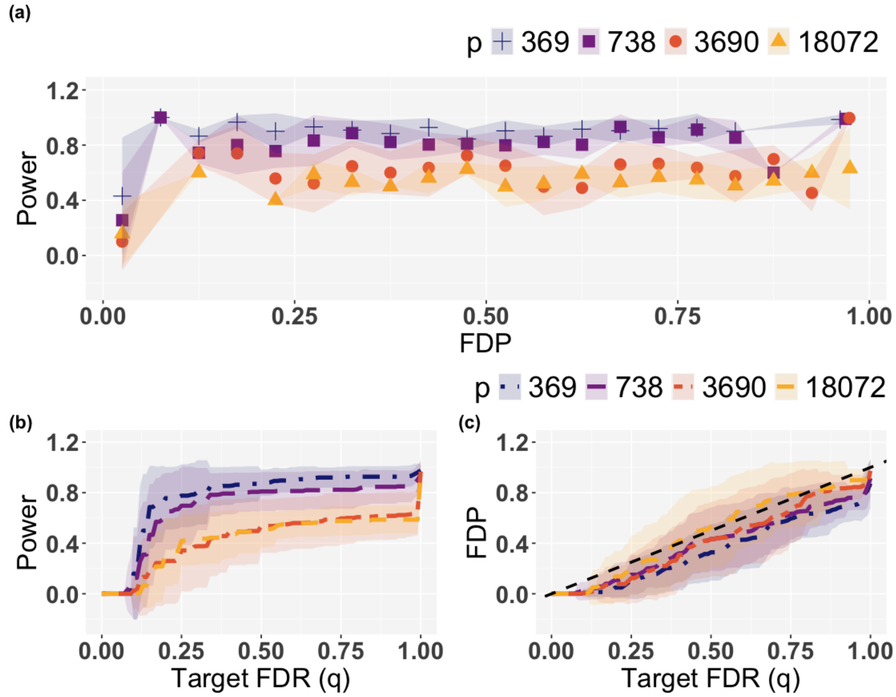

Figure S18: **Comparison of the KO procedure performance for different numbers of null features.** Outcomes are simulated from the CRUKPAP feature matrix with $p \in \{369, 738, 3690, 18072\}$, corresponding to feature-to-sample ratios of 1, 2, 10, and the full feature set, respectively. The linear simulation setting is used with $k = 10$ and $M = 30$ repetitions. (a) Power versus false discovery proportion. Power values are given as means over FDP bins of width 0.05. (b) Power versus target false discovery rate. (c) False discovery proportion versus target false discovery rate.

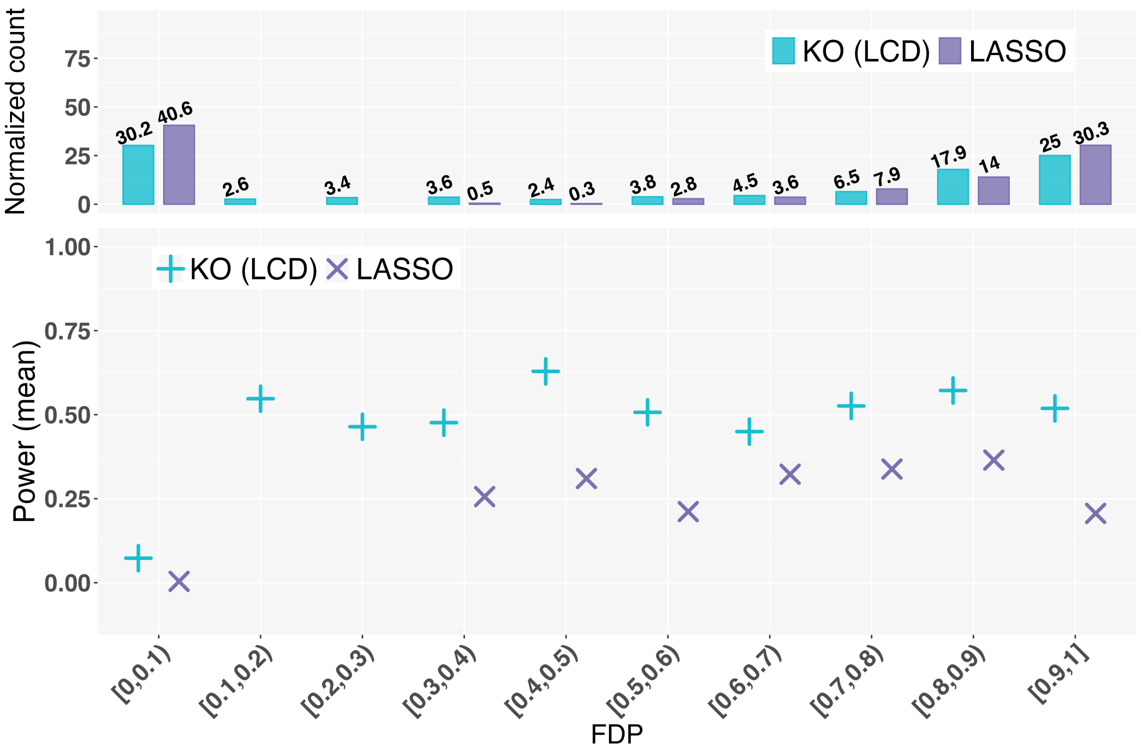

Figure S19: **Comparison of the KO procedure and the lasso penalized logistic regression.** Outcomes are simulated from the complete CRUKPAP feature matrix (369 samples and 18,072 variables) using the linear simulation setting with $k = 10$ and $M = 30$ repetitions. The normalized count gives the number of points in the corresponding FDP interval –*i.e.* the number of points on which the mean power was calculated– normalized by the length of the sequence of parameters for each method (KO: $q$, LASSO: $\lambda$).

### D Supplementary Material: Experiments on additional cohorts

We perform both KO generation methods and statistics comparison with two data matrices built from AEGIS and breast cancer data (see KO generation methods and statistics benchmark for experimental details). For both cohorts we take the intersection between the set of 749 genes used to build $X_{\text{CRUKPAP}}$ and the available gene expression levels in the AEGIS and breast cancer data (see 3.1). For AEGIS all samples are used ($X_{\text{AEGIS}} \in \mathbb{R}^{505\times 727}$) whereas for the breast cancer cohort, in order to remain in a high-dimensional setting, we randomly select 500 samples ($X_{\text{BC}} \in \mathbb{R}^{500\times 736}$).

#### D.1 Figures: KO generation methods and statistics comparison for the AEGIS cohort

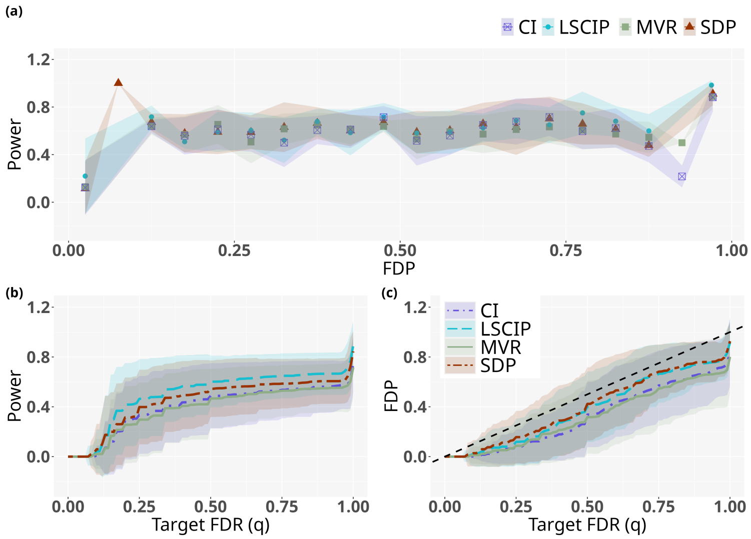

Figure S20: **Comparison of the KO construction methods**. Outcomes are simulated from the AEGIS feature matrix using the linear simulation setting with $k = 10$ and $M = 100$ repetitions. Important variables are selected using the LCD statistics for all KO generation methods. (a) Power versus false discovery proportion. Power values are given as means over FDP bins of width 0.05. (b) Power versus target false discovery rate. (c) False discovery proportion versus target false discovery rate. CI: Conditional independence, LSCIP: Linear Sequential Conditional Independant Pairs algorithm, MVR: Minimum Variance-based reconstruction, SDP: semi-definite programming, see section Generation of KOs features for details.

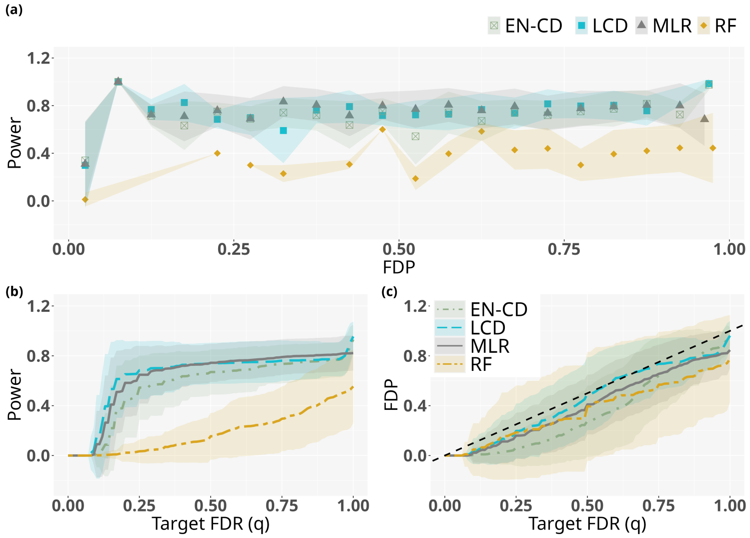

Figure S21: **Comparison of different KO statistics.** Outcomes are simulated from the AEGIS feature matrix using the linear simulation setting with $k = 10$ and $M = 100$ repetitions. KO features are generated using the LSCIP algorithm. (a) Power versus false discovery proportion. Power values are given as means over FDP bins of width 0.05. (b) Power versus target false discovery rate. (c) False discovery proportion versus target false discovery rate. EN-CD: Elastic Net Coefficients Difference, LCD: Lasso Coefficients Difference, MLR: Maximum Likelihood Ratio, RF: Random Forest, see subsection KO test statistics for details.

#### D.2 Figures: KO generation methods and statistics comparison for the BC cohort

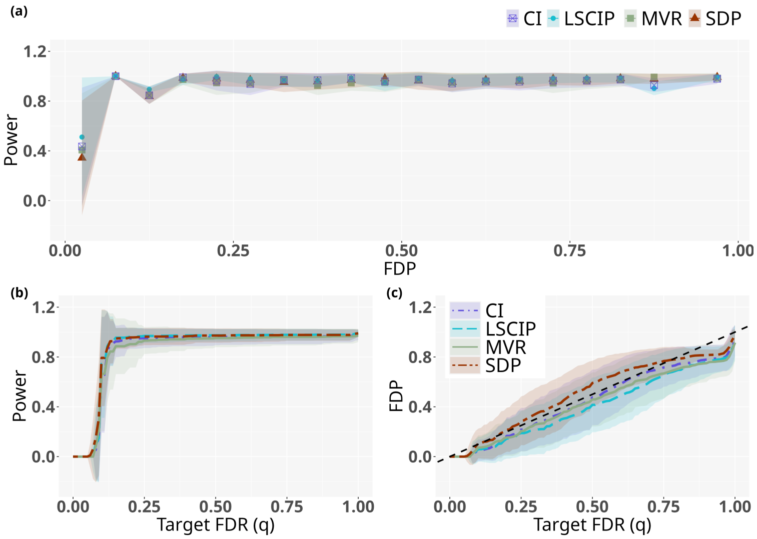

Figure S22: **Comparison of the KO construction methods**. Outcomes are simulated from the breast cancer cohort feature matrix using the linear simulation setting with $k = 10$ and $M = 100$ repetitions. Important variables are selected using the LCD statistics for all KO generation methods. (a) Power versus false discovery proportion. Power values are given as means over FDP bins of width 0.05. (b) Power versus target false discovery rate. (c) False discovery proportion versus target false discovery rate. CI: Conditional independence, LSCIP: Linear Sequential Conditional Independant Pairs algorithm, MVR: Minimum Variance-based reconstruction, SDP: semi-definite programming, see section Generation of KOs features for details.

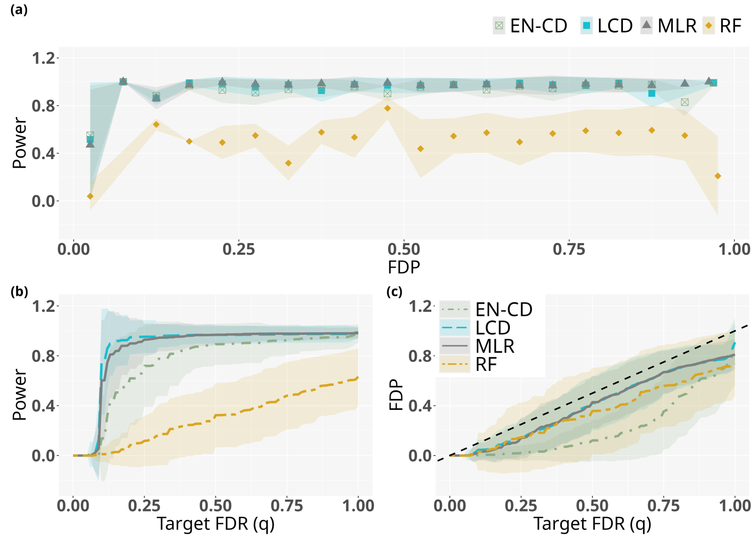

Figure S23: **Comparison of different KO statistics.** Outcomes are simulated from the breast cancer cohort feature matrix using the linear simulation setting with $k = 10$ and $M = 100$ repetitions. KO features are generated using the LSCIP algorithm. (a) Power versus false discovery proportion. Power values are given as means over FDP bins of width 0.05. (b) Power versus target false discovery rate. (c) False discovery proportion versus target false discovery rate. EN-CD: Elastic Net Coefficients Difference, LCD: Lasso Coefficients Difference, MLR: Maximum Likelihood Ratio, RF: Random Forest, see subsection KO test statistics for details.

#### D.3 Comparison KO framework, LASSO, and Wilcoxon rank-sum test with the breast cancer dataset

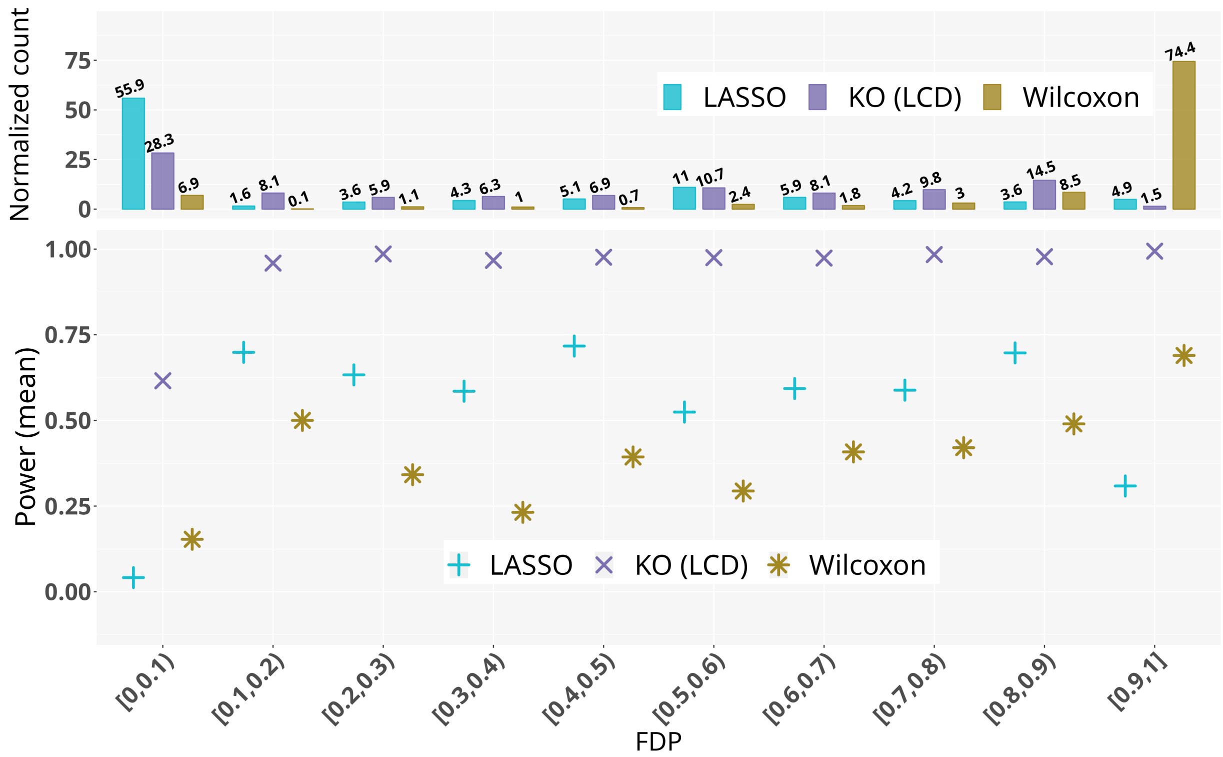

Figure S24: **Comparison of different feature selection methods: the KO procedure, the lasso penalized logistic regression, and the Wilcoxon rank-sum test** Outcomes are simulated from the breast cancer feature matrix using the linear simulation setting with $k = 10$ and $M = 100$ repetitions. The normalized count gives the number of points in the corresponding FDP interval –*i.e.* the number of points on which the mean power was calculated– normalized by the length of the sequence of parameters for each method (KO: $q$, LASSO: $\lambda$, Wilcoxon: $\alpha$) (for details see Subsection Experimental setups).

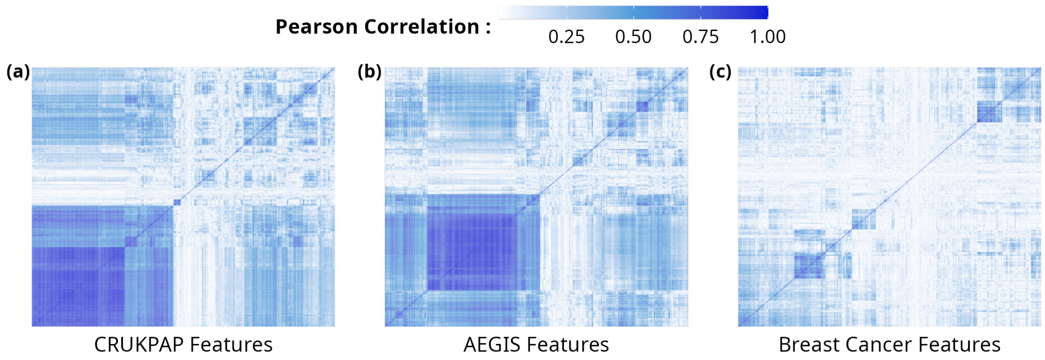

Figure S25: $X_{\text{CRUKPAP}}$, $X_{\text{AEGIS}}$, and $X_{\text{BC}}$ correlation matrices

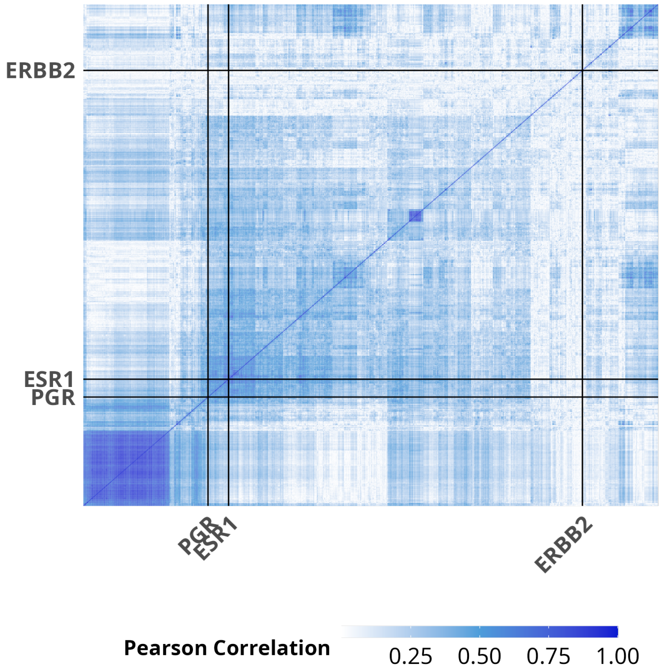

Figure S26: Correlation matrix of the breast cancer data matrix where features correspond to the 869 genes that had non zero weights in at least one of the five biomarkers classifiers [28].

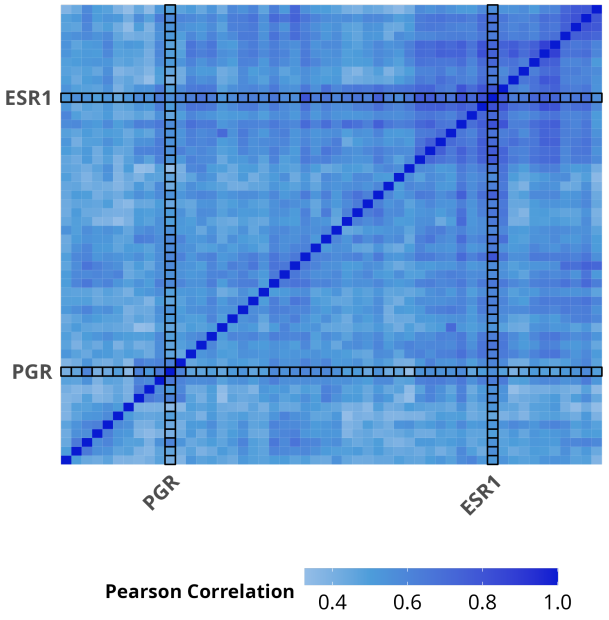

Figure S27: Zoom of the previous correlation matrix of the breast cancer data matrix (see S26 around the ESR1 and PGR genes. Features are among the 869 genes that had non zero weights in at least one of the five biomarkers classifiers [28].

### E Supplementary Material: Correlation study

#### E.1 Figures: Correlation matrices

### F Supplementary Material: Application to real data

| **Cohorts:** | **CRUKPAP** | **AEGIS** | **BC (HER2)** | | **BC (ER)** |
|---|---|---|---|---|---|
| **Target FDR level:** | 0.5 | 0.5 | 0.5 | 0.3 | 0.5 |
| **Nomenclature:** | **Ensembl gene ID** | **Probeset** | **HGNC** | | **HGNC** |
| | ENSG00000002549 | 7893862 | ERBB2 | ERBB2 | ESR1 |
| | ENSG00000170961 | 7894737 | GRB7 | GRB7 | |
| | ENSG00000188385 | 7896908 | MIEN1 | MIEN1 | |
| | | 7985317 | STARD3 | STARD3 | |
| | | 8006504 | PIP5KL1 | PSMD3 | |
| | | 8156058 | AUNIP | PIP5KL1 | |
| | | 7988132 | | SERHL3 | |
| | | | | NTSM | |
| | | | | IRS1 | |
| | | | | STX1A | |
| | | | | AUNIP | |
| | | | | CACNA1D | |
| | | | | RAB17 | |

Table S2: **Genes selection with KOPI for the AEGIS, CRUKPAP and Breast Cancer (BC) data sets**. The selection is made for different values of target false discovery rate (FDR) level. The outcomes are cancer status for AEGIS and CRUKPAP data, and two breast cancer biomarkers status (HER2 and ER), see Real data sets for details.
